# The input-output relation of primary nociceptive neurons is determined by the morphology of the peripheral nociceptive terminals

**DOI:** 10.1101/2020.07.30.228015

**Authors:** Omer Barkai, Rachely Butterman, Ben Katz, Shaya Lev, Alexander M. Binshtok

## Abstract

The output from the peripheral terminals of primary nociceptive neurons, which detect and encode the information regarding noxious stimuli, is crucial in determining pain sensation. The nociceptive terminal endings are morphologically complex structures assembled from multiple branches of different geometry, which converge in a variety of forms to create the terminal tree. The output of a single terminal is defined by the properties of the transducer channels producing the generation potentials and voltage-gated channels, translating the generation potentials into action potential firing. However, in the majority of cases, noxious stimuli activate multiple terminals; thus, the output of the nociceptive neuron is defined by the integration and computation of the inputs of the individual terminals. Here we used a computational model of nociceptive terminal tree to study how the architecture of the terminal tree affects input-output relation of the primary nociceptive neurons. We show that the input-output properties of the nociceptive neurons depend on the length, the axial resistance, and location of individual terminals. Moreover, we show that activation of multiple terminals by capsaicin-like current allows summation of the responses from individual terminals, thus leading to increased nociceptive output. Stimulation of terminals in simulated models of inflammatory or nociceptive hyperexcitability led to a change in the temporal pattern of action potential firing, emphasizing the role of temporal code in conveying key information about changes in nociceptive output in pathological conditions, leading to pain hypersensitivity.

**Significance statement:** Noxious stimuli are detected by terminal endings of the primary nociceptive neurons, which are organized into morphologically complex terminal trees. The information from multiple terminals is integrated along the terminal tree, computing the neuronal output, which propagates towards the CNS, thus shaping the pain sensation. Here we revealed that the structure of the nociceptive terminal tree determines the output of the nociceptive neurons. We show that the integration of noxious information depends on the morphology of the terminal trees and how this integration and, consequently, the neuronal output change under pathological conditions. Our findings help to predict how nociceptive neurons encode noxious stimuli and how this encoding changes in pathological conditions, leading to pain.

## Introduction

Peripheral terminals of nociceptive neurons are key structures in detecting, translating and transmitting noxious stimuli towards the CNS, thus setting the scene for the sensation of pain (Woolf and Ma, 2007; Basbaum et al., 2009; Dubin and Patapoutian, 2010; Gold and Gebhart, 2010; Binshtok, 2011). Single nociceptive terminals combine into complex terminals trees, which differ in number and form of their terminal branches. In some nociceptive terminal trees, the terminals don’t branch after leaving a fiber and terminate at the target organ with a single terminal branch of different length (Zylka et al., 2005; Ivanusic et al., 2013; Alamri et al., 2015; Olson et al., 2017; Alamri et al., 2018; Bouheraoua et al., 2019). Other nociceptive terminals branch within the epithelium, creating a more complex termination (Ivanusic et al., 2013; Alamri et al., 2015; Olson et al., 2017; Alamri et al., 2018). Thus, noxious stimuli activate morphologically complex excitable structures, which differ among primary nociceptive neurons. Importantly, the morphology of terminal trees changes with age or under pathological conditions (Cain et al., 2001; Chartier et al., 2018; Lakatos et al., 2020; Leibovich et al., 2020).

It has been shown that the morphology of dendrites and axons of central neurons (Manor et al., 1991; Mainen and Sejnowski, 1996; Williams and Stuart, 2003; Gidon and Segev, 2012; Jadi et al., 2012; Ferrante et al., 2013) and peripheral mechanosensitive neurons (Lesniak et al., 2014) shape their input-output properties. We, therefore, hypothesized that output of the nociceptive neuron might depend on the integration of outputs from the individual terminal branches and that this integration depends on the structure of the individual terminal and the morphology of the terminal tree.

Noxious stimuli are detected by activation of a specific repertoire of transducer channels expressed in the terminals. The biophysical properties and spatial distribution of functional transducer and voltage-gated channels expressed by nociceptive neurons are essential factors in defining the gain of the terminals, i.e., the number and pattern of action potential firing resulting from activation of the terminal by noxious stimuli (Dib-Hajj et al., 2010; Waxman and Zamponi, 2014; Goldstein et al., 2019). The current electrophysiological and imaging approaches allow studying how a single terminal encodes the information (Vasylyev and Waxman, 2012; Goldstein et al., 2019). Skin-nerve (Reeh, 1986) or teased preparations (Tal and Devor, 1992) permit studying how the whole nociceptive terminal trees respond to stimulation. However, due to technical constraints, it is still unclear what is the contribution of a single terminal location and electrical properties to the output of the terminal tree and how different terminal tree architectures or a change in the specific architecture of the terminal tree relate to the nociceptive function.

Here we implemented a realistic computational model of the nociceptive terminal tree, which we build based on known terminal physiology *in vitro* (Vasylyev and Waxman, 2012) and *in vivo* (Barkai et al., 2017; Goldstein et al., 2019) to study structure-function relation of nociceptive terminals. We stimulated terminals of different terminal trees by a capsaicin-like stimulation and defined the output of the neurons by registering the resulting activity at the central terminal of the modeled nociceptive neuron. We show that the length and the axial resistance of individual terminal branches affect the response of the whole terminal tree to a capsaicin-like current. Our model predicts that the activation of a longer terminal leads to a higher firing rate at the central terminals. Activation of the terminals with lower axial resistance leads to a decreased response. Our model also suggests that the activation of multiple terminals allows summation of responses of each individual terminal, thus facilitating the nociceptive response. Perturbations correlated with inflammatory and neuropathic conditions increase the gain of nociceptive input-output function and change the temporal pattern of the spike trains. Our results show how noxious stimuli are integrated by the nociceptive neurons and predict how structural changes of the nociceptive neurons in pathological conditions can modify nociceptive responses.

## Materials and Methods

Simulations were performed using passive and active properties of the NEURON environment-based compartmental nociceptor model, which we previously developed (Barkai et al., 2017; Goldstein et al., 2019), with adaptations described below. Briefly, we developed a biophysically realistic multi-compartment model of an unmyelinated axon, including the terminal tree (Barkai et al., 2017; Goldstein et al., 2019). The nociceptor morphology consisted of a 25 μm diameter soma-like compartment, connected to a stem axon expanding to peripheral and central axons which join at a T-junction bifurcation site (Figure 1A).

**Figure 1.**
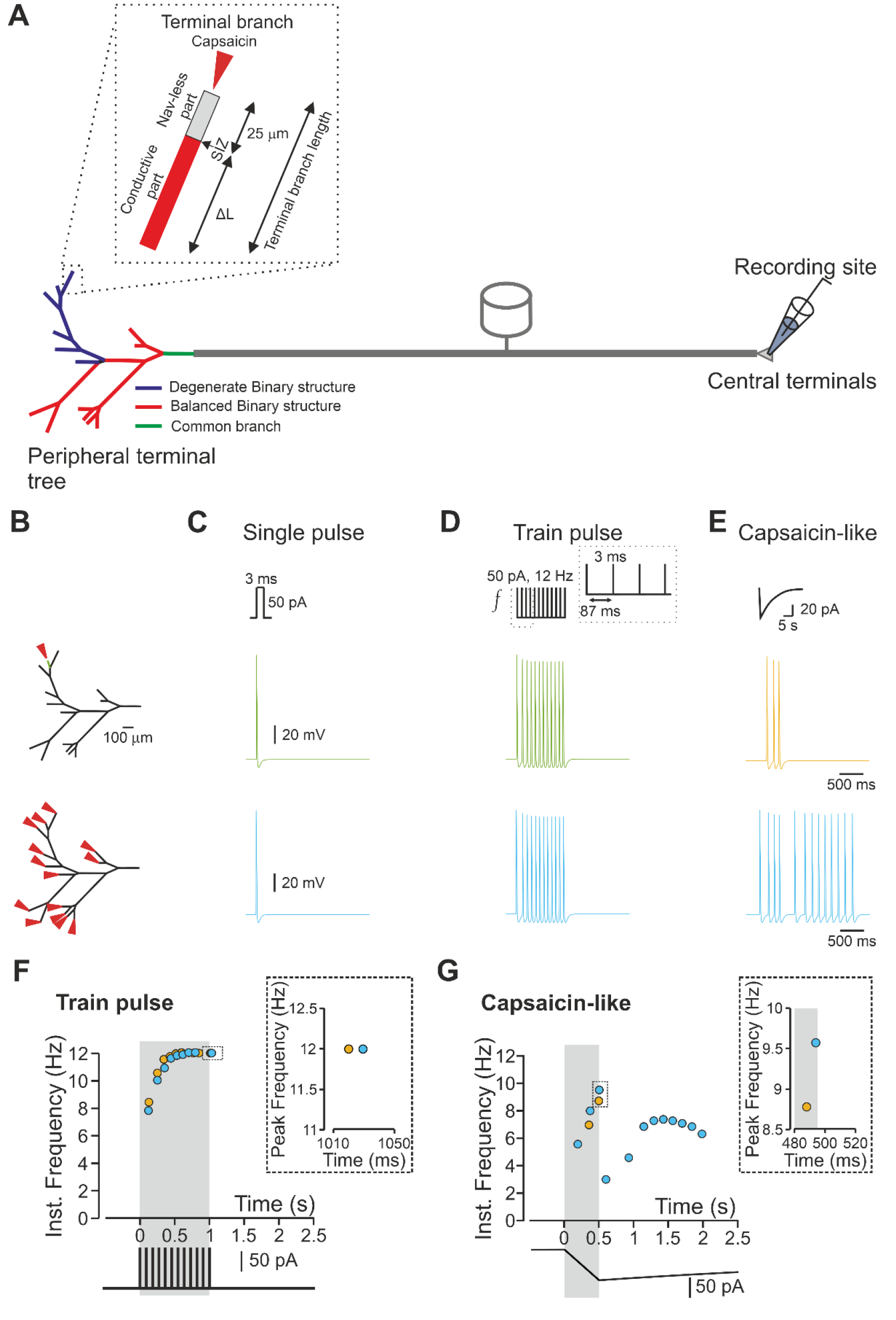
The response of a nociceptive neuron to a capsaicin-like current depends on the morphology of the nociceptive terminal tree. **A.** Scheme depicting the model of the primary nociceptive neuron. The peripheral terminal tree morphology is rendered from the structure of a terminal innervating glabrous skin of mice paw *in vivo*. The terminal tree consists of 14 terminal branches of different lengths. *Inset* depicts the functional organization of a terminal branch. The first 25 μm of each terminal branch does not express Navs (Nav-less, *light grey*). The Nav expression starts from the spike initiation zone (SIZ) and continues for the rest of the fiber (conductive part, *red*). The terminal branch length is defined by the length of the conductive part, which differs among the terminals and the Nav-less part, which has a constant length of 25 μm. The complex terminal tree morphology consists of the simplified version of the balanced binary tree (*red terminal branches*) in which two branches converge into one node (convergence point) and a degenerate binary tree, in which one branch converges into the fiber (*blue terminal branches*). The terminal branches converge into the mother branches, which then converge into the common branch (*green branch*). The stimulations are applied to the terminal branch end (red arrow in the *inset*), and the recording electrode is placed at the central terminal. **B.** The locations of the stimulations. The stimulated terminal or all terminals are marked by the red arrows, and single stimulated terminals are color-coded. **C-D.** Simulated voltage-clamp recordings from a central terminal following stimulation of a single terminal (*yellow*) and all terminals (*light blue*) by single brief 3 ms square current of 50 pA (**C**) or train of brief 3 ms square current of 50 pA applied at 12 Hz (**D**). Note, no difference in firing between stimulating one all terminals. **E.** Same as *C*, but the terminals are stimulated by simulated capsaicin-like current (*see Methods*). Note that in this case, the firing following stimulation of one terminal differs from the firing resulting from stimulation of all terminals. **E.** Instantaneous discharge frequencies following stimulation with a train of brief 3 ms square current of 50 pA applied at 12 Hz (*lower inset*) of a single terminal (*yellow*), and all terminals (*light blue*). Shadowed area outlines the time of the stimulation. Note, no difference in peak instantaneous frequency (*upper inset*). **F.** Same as in *E* but following stimulation with a capsaicin-like current (color-coded as in *E*). Shadowed area outlines the 500 ms of a capsaicin-like current step. Note, an increase in peak instantaneous frequency when all the terminals are stimulated (*upper inset, light blue*).

### Morphology

The morphology of the nociceptive neuron was based on our adaptation of Sundt *et al*.’s model of a nociceptive neuron (Sundt et al., 2015; Barkai et al., 2017). A cylindric soma (25 μm diameter, 25 μm length) was connected to a stem compartment (0.8 μm diameter, 75 μm length) which bifurcate to a T-junction, creating two branches, the first branch (0.8 μm diameter, 100 μm length) connected to a peripheral axon (0.8 μm diameter, 5 mm length) and the second branch (0.4 μm diameter, 100 μm length) connected to a central axon (0.4 μm, 5 mm length). All of the abovementioned compartments, apart from the soma, were subdivided into 100 segments each. The peripheral axon was then connected to a terminal tree structure, as mentioned below.

### Terminal tree structures

For the *“realistic” type of the terminal tree,* which was reconstructed according to one of our arbitrary experimental observations (Barkai et al., 2017), the distal axon ending tapered at its end and expanded into a terminal tree composed of 27 branches. For proper electrical propagation from the terminal tree to the peripheral axon, these two compartments were connected by a tapered cone-like axon with a linearly changing diameter over 100 μm of length.

The branches of the terminal tree were of variable lengths (50 - 300 μm) and of a diameter of 0.25 μm. These branches included 13 mother branches, which were connected from the both ends to other branches and terminal branches (*see below*), which were connected at the proximal end to a mother branch and had a free-nerve ending on the other end (Figure 1A).

#### Binary tree models

Balanced and degenerate binary tree structures were generated in and simulated in PyNeuron. All terminal tree branches, unless otherwise mentioned, were identical in length (50 μm). Binary tree structures were changed according to the number of their bifurcation stages (n_*l*_), which controlled the total number of the terminal branches (n_*TB*_) according to n_*TB*_ = 2^n*l*-1^ (Jarvis et al., 2018).

*The terminal branches* of all types consisted two sections separated by the spike initiation zone (SIZ, Goldstein et al., 2019): a 25 μm long compartment which consisted all the active conductances described below, but was absent of voltage-gated sodium channel (Nav) conductances (Nav-less compartment) and the “conductive” compartment connecting between the Nav-less compartment to the junction with the mother branch (Figure 1A). The conductive compartment possessed all the active conductances described below. In some experiments, the length of either Nav-less or conductive compartments on all their properties was changed.

The 25 μm Nav-less compartment of the terminal branches was sub-divided into 2.5 μm segments (10 segments). The conductive compartment was subdivided into 5 μm segments; thus, the number of segments changed according to the length of the compartment.

### Passive membrane properties

Membrane capacitance of 1μF cm^−2^ was set for all compartments. Membrane resistance of 10000 Ω cm^−2^ was set for all compartments apart from the terminal branch, which has a 4-fold somatic membrane resistance (Vasylyev and Waxman, 2012). Axial resistance (Ra) in all compartments, apart from the terminal branch, was 150 Ω cm^−2^ (Barkai et al., 2017). The axial resistance of the terminal branch (Ra_*TB*_) was set to 15-fold of the Ra (Ra_*TB*_ = 2.25 MΩ cm^−2^). In some experiments, the value of Ra of the conductive part of the terminal branch was changed, and its effect on neuronal excitability was examined.

The passive conductance equilibrium potential was set to −60 mV (Gudes et al., 2015). The simulated resting potential was −58 mV, in line with our experimental results (Barkai et al., 2017).

### Active conductances

The model neuron of all tree structures includes the following Hodgkin-Huxley-type ion channels used in Barkai *et al.* (Barkai et al., 2017) and Goldstein *et al.* (Goldstein et al., 2019): TTX-sensitive sodium current (I_NattxS_), TTX-s sensitive persistent sodium current (I_NaP_), Nav1.9 TTX-r sodium channels (I_Nav1.9_) and Nav1.8 TTX-r sodium channels (I_Nav1.8_). All channels parameters of the sodium currents were adapted from Herzog *et al*. (Herzog et al., 2001) and Baker *et al.* (Baker, 2005). Three types of potassium channels included: (i) the delayed rectifier channel (I_KDR_) adapted from (Herzog et al., 2001); (ii) An A-type potassium channel (I_KA_) adapted from Miyasho *et al.* (Miyasho et al., 2001), whose activation and inactivation gates were shifted by 20 mV in hyperpolarized direction to closely resemble kinetics of DRG neurons (Qu and Caterina, 2016) and (iii) the Kv7/M channels which were adapted from Shah *et al*. (Shah et al., 2008) and tuned to our experimental results. The h-current (*I*_h_) was also included and taken from Shah *et al.* (Shah et al., 2008), the slope factor was tuned according to Komagiri and Kitamura (Komagiri and Kitamura, 2003).

For simulating the excitable properties of a single-compartment neuron we used the following fixed maximal conductance parameters:

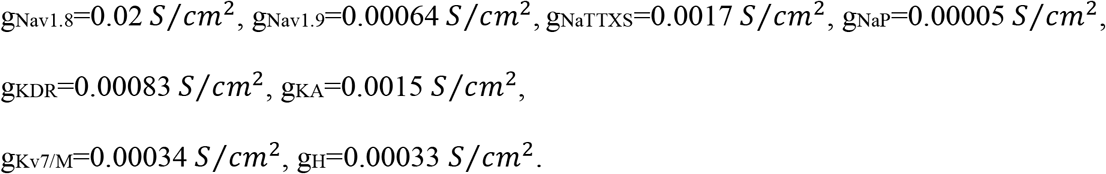

Calcium channels were adapted directly from Shah *et. al.* (Shah et al., 2008) and their conductance values were tuned according to the inward currents described in Blair and Bean (Blair and Bean, 2002). The T-type and L-type channels representing the low voltage-activated (LVA) and high voltage-activated (HVA) currents, respectively, were added with the following conductance values:

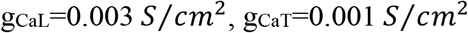

The reversal potentials for sodium (E_Na_), potassium (E_K_), and h-current (E_H_) were set to +60mV, −85mV and −20 mV, respectively. The passive reversal potential was set to −60mV.

Apart from Nav conductances, which were unevenly distributed between Nav-less and conductive compartments, all other conductances were evenly distributed in all compartments.

*The recordings* were performed by positioning a NEURON “point process” electrode at the terminal end of the central axon (Figure 1A). In some experiments, the recording electrode was positioned at the Nav-less compartment, 2.5 mm before the SIZ or at the conductive compartment of the terminal branch, 2.5 mm after the SIZ.

### Stimulation parameters

Simulations were performed, assuming a room temperature of 25° C. To avoid boundary condition problems, the stimulating electrode or capsaicin puff-like process were positioned at 10% of the terminal branch length taken from the terminal branch’s tip.

For simulating an optogenetic stimulation, a short 3 ms square pulse of either 50 or 100 pA was applied into a single simplified voltage-clamp point-process at the distal end of the single or multiple terminal branches. In some experiments, these brief square pulses were applied at various frequencies.

Capsaicin-like stimulation was injected to single or multiple nerve-endings. A capsaicin-like induced current was introduced into a single simplified voltage-clamp point-process with fast exponential activation and slow exponential inactivation mimicking the experimental kinetics of puff-applied 1 μM capsaicin-induced current and sufficient to induce action potential firing when applied onto the acutely dissociated dorsal root ganglion neuron (Nita et al., 2016). The similar current was used in Barkai *et al.* (Barkai et al., 2017) and Goldstein *et al.* (Goldstein et al., 2019) to generate activity in the modeled nociceptive neuron when applied to the single 75 μm length terminal:
for *t* ≤ *t_puff_*

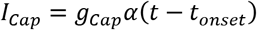

for *t_puff_* < *t*

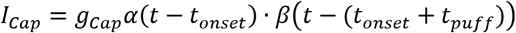

where *g_Cap_* is the maximal conductance, *α(x)* and *β(x)* are the activation and inactivation functions with *τ_α_* and *τ_β_* as the activation and inactivation time constants respectively and were applied with the following values:

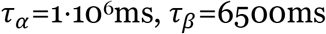

*t_onset_* and *t_puff_* are the times at which the puff application simulation begins and the length of application, respectively.

To mimic the changes in the membrane resistance following the opening of TRPV1 channels, the capsaicin-like current injection was accompanied by an opening of passive transducer channels (Goldstein et al., 2019). The conductance to this channel 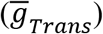 was exponentially distributed and the exponential decay constant (*γ)* was calculated to fit the diffusion of capsaicin and hence change in its concentration as a function of distance from the pipette tip as was previously calculated in Goldstein *et. al.* (Goldstein et al., 2017). In the experiments where the length of the terminal branches was changed, the distribution of the transducer conductances was adapted accordingly.

The final transducer distribution equation introduced into the model was:

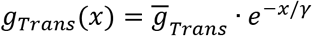

Where *x* is the distance from the nerve-ending and the fixed parameters were:

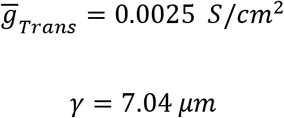

### Randomized onset timing

Onset time was generated with an addition of a normal distribution randomize value *N_R_*:

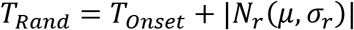

Where *μ* is the mean (and set to zero) and *σ_r_* is the standard deviation. We modify the value of *σ_r_* to generate randomized onset timing. The randomized part of the equation was an absolute value since the onset of stimulation cannot occur prior to the stimulation time. Each generated stimulation was then applied to a single terminal.

### “Inflammation” (SIZ shift) model

To simulated inflammatory hyperalgesia, we mimicked the distal shift of the SIZ (Goldstein et al., 2017) by shortening of the Nav-less compartment from its normal 25 μm lengths to 20 μm. The terminal’s conducting compartment was elongated by the same length reduced from the passive compartment, leaving the total terminal length at a constant value. All the active and passive parameters were modified accordingly.

### “Neuropathic pain” (Noise) model

To simulated nerve injury mediated hyperalgesia, an external noise conductance was incorporated into all terminal tree branches as previously described (Olivares et al., 2015, Barkai et al. 2017 et al.). The noise was injected into the whole terminal tree and was based on the Ornstein-Uhlenbeck process with mean 0, and the current was taken from Olivares et al. (Olivares et al., 2015).

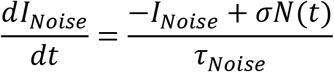

Where σ is a square root of the steady-state variance of the current amplitude; N - is a normally distributed random variable with zero mean and variance = 1 and τ is the steady-state correlation time length of the current (was fixed to 1 ms).

### Data Analysis

Unless otherwise stated, the firing patterns were recorded from the central terminal compartment. The first 500 ms were of each recording were discarded in the analysis in order to remove possible transient firing patterns.

Instantaneous spike frequencies were analyzed and plotted as previously described by Zimmerman et al. (Zimmermann et al., 2009).

### Data and software availability

All datasets generated during and/or analyzed during the current study are available in the main text or upon request from the Corresponding Author, Alexander Binshtok.

The model files will be uploaded to Model DB and also available upon request from the Corresponding Author, Alexander Binshtok.

## Results

Nociceptive terminal trees are complex structures composed of individual branches of different lengths and complexity (Treede et al., 1990; Zylka et al., 2005; Ivanusic et al., 2013; Alamri et al., 2015; Olson et al., 2017; Alamri et al., 2018; Bouheraoua et al., 2019). The terminal branches, which ends at the target organ and detect the noxious stimuli consist of (1) the terminal, which is activated by noxious stimuli; (2) the passive part, a first 20-30 μm of the terminal branch, in which the depolarization, resulting from the terminal activation, propagates without involving voltage-gated sodium channels (Navs) and (3) the conductive part where propagation is active as it involves Navs (Goldstein et al., 2019). The length of the terminal and the passive part of the terminal branch is similar among different terminals (Goldstein et al., 2019); thus, the length of the conductive part defines the overall length of the terminal branch. The terminal branches converging onto the fibers constructing the nociceptive terminal tree (Belmonte et al., 2004). To examine the effect of the morphology of the nociceptive terminal trees on their coding properties, we adapted a computational model of the nociceptive nerve fiber, which we previously described (Barkai et al., 2017; Goldstein et al., 2019). In this model, we first based the geometry of the terminal tree on the rendering of a nociceptive terminal tree visualized using *in vivo* two-photon microscopy (Barkai et al., 2017). The complex structure of the “realistic” modeled terminal tree can be simplified and represented as a combination of different “tree data structures”: a balanced binary tree, where each branching point, a node, bifurcates into two branches (Figure 1A, *red branches*) and a degenerate binary tree, where each node give rise only to one branch (Figure 1A, *blue branches*). The nodes are branching points from an anatomical point of view. However, from a physiological point of view, these nodes in afferent neurons serve as merging or convergence points of voltage deflections from the converging terminal branches.

We adapted the physiological properties of the terminals from data obtained from recordings of capsaicin-mediated activity of corneal nociceptive terminals *in vivo* (Goldstein et al., 2019). Specifically, the contribution of Navs began only 25 μm proximal to the terminal tip (Nav-less compartment, Figure 1A, *inset*); thus, the spike initiation zone is located 25 μm from the tip (Goldstein et al., 2019). Also, we increased the axial resistance of the terminal branches to reflect the presence of intracellular organelles, such as mitochondria (Heppelmann et al., 1994; Müller et al., 1996; Heppelmann et al., 2001; Alamri et al., 2018). In these conditions, the action potential (AP) conduction velocity along the terminal branch was 0.04 m/s, and the refractory period for a 3 ms, 50 pA stimuli was 49 ms. The refractory period in nociceptive c-fibers, obtained by stimulating the central process and recording from cell somata, was shown to vary between 4 to 45 ms (Gemes et al., 2013).

We first compared the nociceptive output following activation of a single terminal activation of all the terminals in the “realistic” terminal tree to examine how the structures of terminal trees affect the gain of the nociceptive neurons, i.e., the relation between the stimuli and the resulted AP firing (Kispersky et al., 2012). We activated the terminals with simulated rectangular or capsaicin-like currents (*see Methods*) and monitored the resulting activity by measuring the number of APs and their instantaneous frequency at the central terminal (*see Methods*, Figure 1A). The stimulation was applied at the tip of a random terminal or all the terminals (Figure 1B, red arrowhead; *see Methods*). First, we simulated the activation of a single terminal (Figure 1B, C, *green*) with a length of 75 μm (25 μm of the Nav-less compartment + 50 μm conductive compartment, Figure 1A, *inset*) using a brief (3 ms) suprathreshold (50 pA) rectangular current. We chose these parameters since the optogenetic stimulation of a similar duration was sufficient to generate a single AP (Browne et al., 2017). Indeed, the stimulation of this terminal with a single short current pulse produced a single AP at the central terminal (Figure 1C, *upper panel*, *green*). We next simulated the state in which the stimulus affects a wider area and simultaneously activates all terminals in the same receptive field (Treede et al., 1990). We asked whether stimulation of multiple terminals would amplify the resulted response of a nociceptive neuron. We simultaneously stimulated all terminals from the same fiber by a brief suprathreshold stimulus and showed that still only one AP was recorded at the central terminal (Figure 1B, C, *lower panel, light blue*). Train of short single-action potential-generating stimuli applied to a single terminal at 12 Hz produced a train of APs at the central terminals with identical frequency (Figure 1D, *upper panel, green)*. Moreover, the application of brief multiple stimuli to all terminals (Figure 1D, *lower panel, light blue*) did not produce any increase in the number of APs fired. The simultaneous stimulation of all terminals did not improve the firing frequency (Figure 1F). The increase in the stimulus amplitude to 100 pA also did not change the output of the terminals such that the modeled cell fired 12 APs for a 12 Hz stimulation (*data not shown*). These data suggest that activation of a single terminal by brief stimuli conveys the information similar to the information integrated from multiple terminals.

However, we previously showed that stimulation of a simulated single terminal by prolonged continuous capsaicin-like current leads to firing of up to five APs (Goldstein et al., 2019). Recordings from C and Aδ fibers in skin-nerve preparation or teased fiber preparation showed that application of noxious stimuli to a receptive field, i.e., activation of multiple terminals, lead to a much higher AP firing (Levy et al., 2000; St Pierre et al., 2009; Zimmermann et al., 2009; Murthy et al., 2018; Vandewauw et al., 2018). These results, which describe the activation of the whole receptive field by a prolonged “natural” stimuli, suggesting that activation of multiple terminals by prolonged stimuli may lead to a summation of the responses and overall increase in AP number. To examine this hypothesis, we stimulated either a single terminal or all the terminals by a simulated capsaicin-like current (Figure 1E, *see Methods*) to mimic the “natural” activation of TRPV1 channels located at the terminals (Goldstein et al., 2019). The simulated capsaicin current (Figure 1E) was modeled from a current evoked following a 500 ms puff application of 500 nM capsaicin, recorded from acutely dissociated nociceptive DRG neurons, held at −65 mV (Nita et al., 2016; Barkai et al., 2017; Goldstein et al., 2019). Stimulation of a single 75 μm terminal with a capsaicin-like current triggered 3 APs at the central terminal (Figure 1E, *upper panel, green*). Importantly, stimulation of all terminal branches with the same capsaicin-like current substantially increased the AP firing (13 Aps, Figure 1E, *lower panel, light blue*), amplified the firing frequency (Figure 1F), and shorten the first spike latency (191 ms when all terminals are stimulated vs. 345 ms when only single terminal was stimulated).

These data predict that stimulation of multiple terminals by prolonged capsaicin-like current facilitates the response by increasing the firing and response frequency, differently from the activation by the brief pulses. This implies that wider activation of the terminal tree will increase the nociceptive response.

The activation of all terminals introduces a number of variables that may underlie the increased nociceptive output: (1) the increase in firing following stimulation of all terminals may result from the integration and the summation of the responses from individual terminals at the convergence points; (2) since the structure of the tree is asymmetrical, the distance of the individual terminal from the common convergence point is different; thus the effect of the closest terminal may affect the overall nociceptive response; and (3) stimulation of all terminals activates terminals with different electrical properties, e.g., different length of the conductive part, which may contribute to the difference in the activation of a single terminal to that of multiple terminals. We, therefore, systematically studied the effect of these factors on nociceptive gain.

To prevent the effect of asymmetry in length and in the terminal location on nociceptive excitability and to examine only the effect of a number of activated terminals, we studied the impact of multiple stimulations on the symmetrical (balanced) binary trees (Figure 1A) with similar terminal branch lengths. First, we applied a capsaicin-like current to either one or both terminal branches of a simple, symmetrical, balanced binary tree containing only one node and two symmetrical branches (Figure 2A, *upper panel*). We measured the resulted changes in terminal voltage after the convergence point. Stimulation of a single terminal resulted in a single AP (Figure 2A, *bottom*, *blue trace*). Simultaneous stimulation of both terminals by a capsaicin-like current generated two APs (Figure 2A, *bottom*, *red trace*). This result is unexpected, as we assumed that at the convergence point, only one AP would propagate, resetting the other branches (Weidner et al., 2003; Gemes et al., 2013). We hypothesized that stimulation with prolonged capsaicin-like current, differently from stimulation with a brief square-like current, might induce a long-lasting after-AP depolarization that persists after a refractory period of the first AP is over. In a case of a single stimulus, this after-AP depolarization may not be sufficient to generate additional AP. Simultaneous stimulation of both terminals might lead to the summation of after-AP depolarization from two terminal branches, which, in this case, maybe suprathreshold and sufficient to generate a second AP. To examine this hypothesis and to examine if, indeed, simultaneous stimulation facilitates after AP depolarization, we measured the voltage deflection resulting from a single stimulus or simultaneous stimulation of two terminals. To prevent the effect of voltage-gated sodium channels on the voltage deflection, we removed all Nav conductances from the simulation. Simultaneous stimulation of two terminals did not affect the voltage deflection measured at the SIZ of each of the branches (Figure 2B) but substantially increased the voltage deflection after the convergence point (Figure 2C). The summated voltage deflection persisted (Figure 2C, *grey shadowed area*), succeeding refractory period of the AP (49 ms, Figure 2C, *red shadowed area*). These data suggest that an increase in the number of activated terminals facilitates nociceptive activation, by allowing the summation of the depolarizations from the converging terminal branches. To further validate this conclusion, we increased the complexity of the balanced binary tree type of the nociceptive terminal (Figure 3). We systematically examined the activation of a single terminal branch, the simultaneous stimulation of two adjacent (sisters) terminal branches and all terminal branches at the terminal trees composed of a different number of terminal branches (n_*TB*_) defined by the number of the “bifurcation stages” (n_*l*_) according to n_*TB*_ = 2^n*l*-1^ (Figure 3A, *see Methods*, Jarvis et al., 2018). Activation of a single terminal branch by a capsaicin-like current in n_*l*_ = 2 tree with two terminals, evoked one AP at the central terminal (Figure 3C, Figure 2) and two APs in more complex examined trees (Figure 3B, C, *yellow*). As our model predicts (Figure 2), activation of two (sister) terminals led to a moderate increase in firing at the central terminals (Figure 3B, C, *purple*).

**Figure 2.**
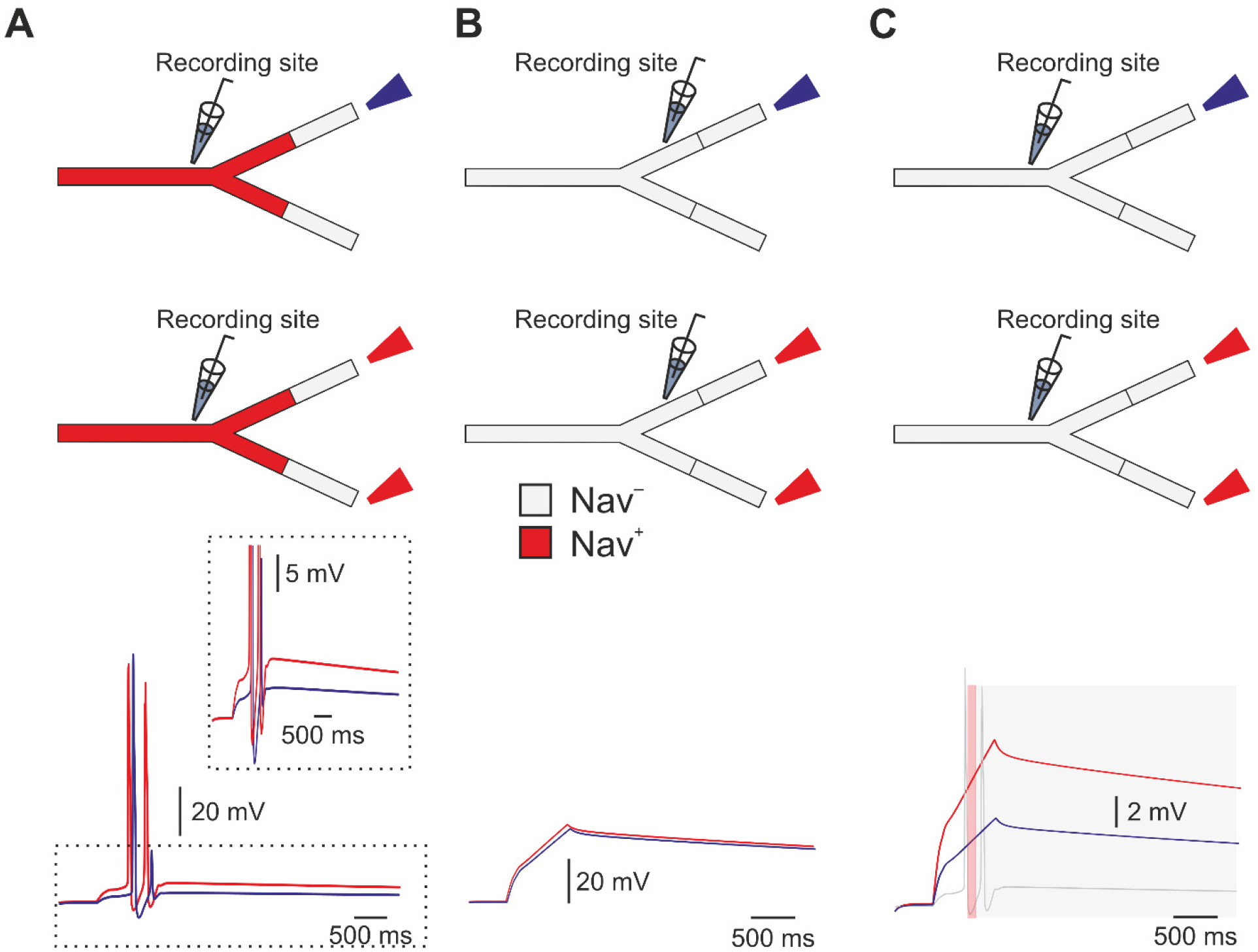
Stimulation of multiple terminals allows summation of the voltage and facilitates nociceptive response. **A.** *Upper*, scheme depicting the experimental conditions: either one (*blue*) or both (*red*) 75 μm length terminal branches converging into the same convergence point in the symmetrical binary tree were stimulated by a capsaicin-like current. *Lower*, the resulted response recorded after the convergence point. Note that simultaneous activation of both terminals generated two APs (red trace), with higher after depolarization (*inset*). **B.** With all the Nav conductances annulled, activation of either one or two terminals with a capsaicin-like current produced similar depolarization measured before the bifurcation **C.** Activation of two terminals led to substantially higher depolarization (*red*) after the converging point than activation of only one terminal (*blue*). The AP firing following activation of two terminals, shown in *A* is superimposed with the voltage traces to show that the depolarization evoked by the activation of two terminals remained high also at the time after the refractory period of the AP. The red shadowed area outlines the refractory period (49 ms), and the grey shadowed area outlines the time of the summated post-AP depolarization.

**Figure 3.**
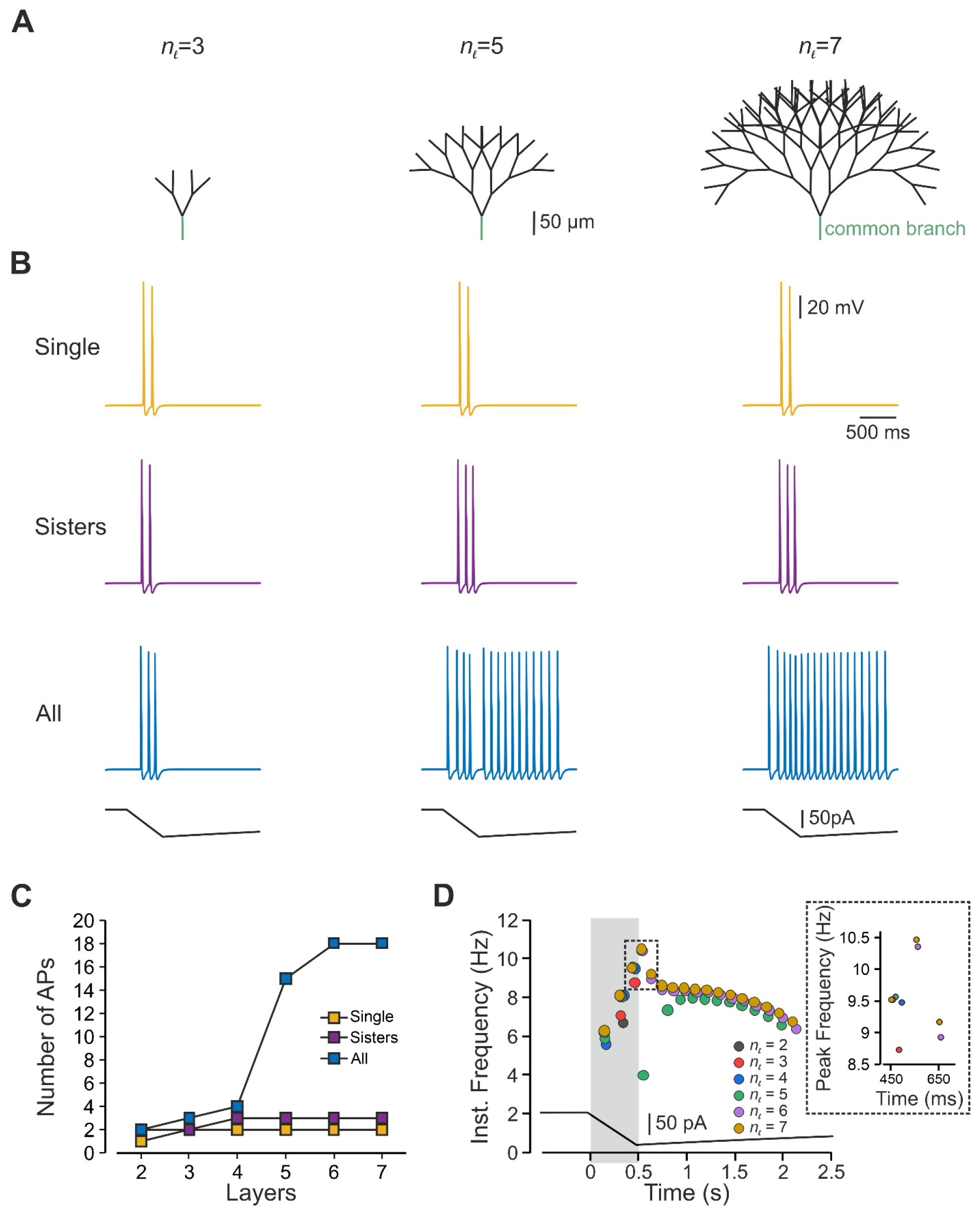
In a symmetrical nociceptive terminal tree, an increased number of stimulated terminals facilitates nociceptive response. **A.** Scheme depicting the model of the symmetrical binary trees of different complexities determined by the number of bifurcation stages (n*_l_*, *see Methods*), which defines the number of terminal branches. **B.** Responses detected at the central terminal following stimulation of either single terminal branch (*yellow*), pair of sister terminal branches converging into the same converging point (*purple*) and all terminals (*blue*) by a capsaicin-like current (depicted below the traces) applied to the representative n_*l*_ = 3, n_*l*_ =5 and n_*l*_ = 7 trees. **C.** The number of APs plotted as a function of the number of bifurcation stages, following stimulation of a single terminal branch (*yellow*), pair of sister terminal branches converging into the same convergence point (*purple*) and all terminals (*blue*) by a capsaicin-like current. **D.** Instantaneous discharge frequencies following stimulation with a capsaicin-like current (*shown below*) applied to all terminals of the trees of different complexity (n_*l*_). *Inset*: peak instantaneous frequency (color-coded as in *D*).

Activation of all terminals led to a substantial increase in the number of AP than activation of a single terminal in all examined trees (Figure 3B, *blue*). Importantly, as the complexity increased, i.e., the number of the terminal branched increased, activation of all terminals led to a rise in AP firing at the central terminal (Figure 3B, C, *blue*) with increased firing frequency (Figure 3D). It is noteworthy that although the number of AP changes considerably between stimulations of the terminal trees of different complexity, the instantaneous frequency and peak instantaneous frequency change very little (Figure 3D). Altogether the results from the symmetrical terminal tree analysis suggest that stimulation of more terminals (activation of a larger area or area with higher terminal density) will lead to increased response. It also implies that an increase in the terminal complexity facilitates nociceptive gain.

From a mechanistic point of view, the increase in gain in terminal trees with more bifurcation stages could be due to, as our model predicts, the depolarization summation at the convergence points (Figure 2) or an increase in the length of Nav expressing compartments of the terminal tree, which elongates as the tree complexity increases (Figure 3). The elongation of Nav expressing components was shown to increase neuronal excitability (Kuba et al., 2006). To distinguish between the effect of the activation of multiple branches and that of the length on nociceptive gain, we simultaneously stimulated all four terminals of n_*l*_ =3 tree while changing the length of the common branch (Figure 3A, *green branch*, Figure 4A, *green dashed line*). In our model, the common branch is a part of the terminal tree, accordingly, it possesses all the electrical properties of the mother branches and has a diameter of 0.25 μm. The common branch is connected to the thicker fiber (0.8 μm diameter) by a tapered cone-like axon with a linearly changing diameter. Our data show that an increase in the length of the conductive components indeed leads to an increase in firing and frequency, at the central terminal (Figure 4B), suggesting that the increase in overall length of Nav expressing compartment of the terminal tree plays a role in the increased gain at the more complex terminals. However, the increase in length could account only for part of the increase in gain as stimulation of all terminals in a n_*l*_ =3 tree with a length equivalent to the length of a n_*l*_ = 7 tree (350 μm), although led to a more firing than that of the shorter a n_*l*_ = 3 tree, generated less firing than activation of all terminals in a n_*l*_ = 7 tree of the same length (11 *vs.* 18 APs, Figure 4B, *left*; Figure 3C). Altogether our data predict that activation of more terminals will lead to an increase in AP firing at the central terminal due to the summation of the depolarization at the convergence points, which in case of multiple terminal activations would “meet” higher Nav conductance, leading to more firing.

**Figure 4.**
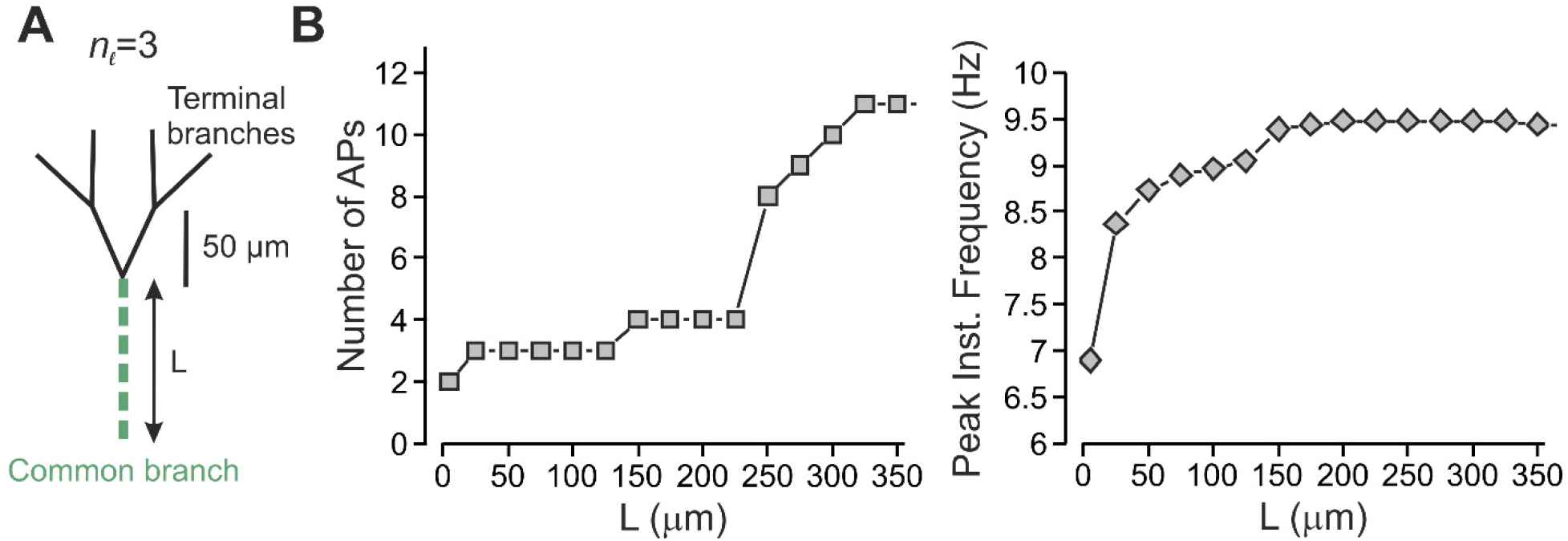
Complex terminal trees affect nociceptive gain partially by virtue of the length of their conductive parts. **A.** Scheme of the experiment, the length of the common branch (*green*) was changed without changing the number of terminal branches. All terminals were simultaneously stimulated by a capsaicin-like current, and the resulted activity was measured at the central terminal. **B.** The number of APs recorded at the central terminal (*left*) and their peak instantaneous frequency (*right*) plotted as a function of the length of the common branch.

In addition to the balanced binary tree, the “realistic” terminal tree also consists of a part that can be simplified to a degenerate binary type of terminal (Figure 1A, *blue branches*). In this type of binary tree, each node gives rise only to one terminal branch of a similar length (Figure 5A). We examined the impact of the complexity of this type of terminal structure on its encoding properties. We used a degenerate tree with branches of a similar length (Figure 5A), thus avoiding possible effects of the length of the terminal branch on the gain. Activation of all terminals in the degenerate tree led to a substantially higher number of APs than activation of a single terminal (Figure 5A, *yellow terminal*), or sister terminals (Figure 5A, *yellow and purple terminals*), in all examined trees (Figure 5B). Also, in terminals with more bifurcation stages, the activation of all terminals led to increased AP firing at the central terminal (Figure 5B, *light blue*), with increased firing frequency (Figure 5B - D). These data suggest that in the degenerate terminal tree activation of more terminals facilitates the response, similar to the balanced binary type of terminal tree. However, here, in addition to the number of the convergence points, and the overall length of the Nav containing components, an additional level of complexity exists. In the balanced binary tree, the distances between each single terminal end to the beginning of the common branch are equal (Figure 3A) such that the generated responses would arrive at the convergence points at the same time. In the degenerate binary tree, however, the distance between the different terminals to the common branch (Figure 5A, *green branch*) are different (for example, the difference at a distance to a common branch between yellow and light blue terminals in Figure 5A, n_*l*_ = 7 tree). This suggests that the signals from the simultaneously activated terminals would arrive at the common branch at different times and if one of the branches is situated closer to the common convergence point, the APs generated at this branch would reach the common convergence point before the APs generated by other terminals which are situated further away. The AP generated at the closest terminal branch would propagate both orthodromically and antidromically, upon reaching the convergence point. In the latter case, it would render the rest of the terminal tree inactive, by inactivating Navs, and if the firing and following inactivation are long enough, it may prevent APs from other terminal branches to reach the central terminal. We stimulated the closest terminal to the common branching point of the “realistic” terminal tree with a capsaicin-like current to examine if, indeed, the stimulation of the branch closest to the common convergence point would take over the activation of the whole terminal tree (Figure 6A, *deep navy-blue terminal*). This terminal was of 175 μm length, and its activation led to the firing of 13 APs, which indeed reached the central terminal faster than other terminals (191 ms vs. for example 345 ms from the terminal shown in Figure 1A, *bottom*) and at the same time as when all the terminals were active (191 ms, both). Moreover, the activation of the closest terminal led to an identical pattern of firing to the firing evoked by stimulation of all terminals (Figure 6A *right* and B). These data suggest that the location of the terminal relative to the common convergence point is important in defining the effect of a terminal on the nociceptive firing. We next examined how activation of other terminals contributes to nociceptive activity when the closest terminal does not produce dominant firing. The shortening of the closest terminal to 75 μm evoked three APs following activation by a capsaicin-like current (Figure 6C, *deep navy blue*), and activation of sister terminals evoked four APs (Figure 6C, *green*). In this case, activation of all terminals produced the firing of 13 APs (Figure 6, *dark green*), leading to the facilitation of nociceptive response. The results of this experiment predict that the location of the individual terminal is important in defining the overall contribution of this terminal to the nociceptive response. The results showing that activation of the terminals of different length (75 vs. 175 μm) evoke different firing also suggest that apart of the location, the electrical properties of the individual terminals affect the nociceptive gain.

**Figure 5.**
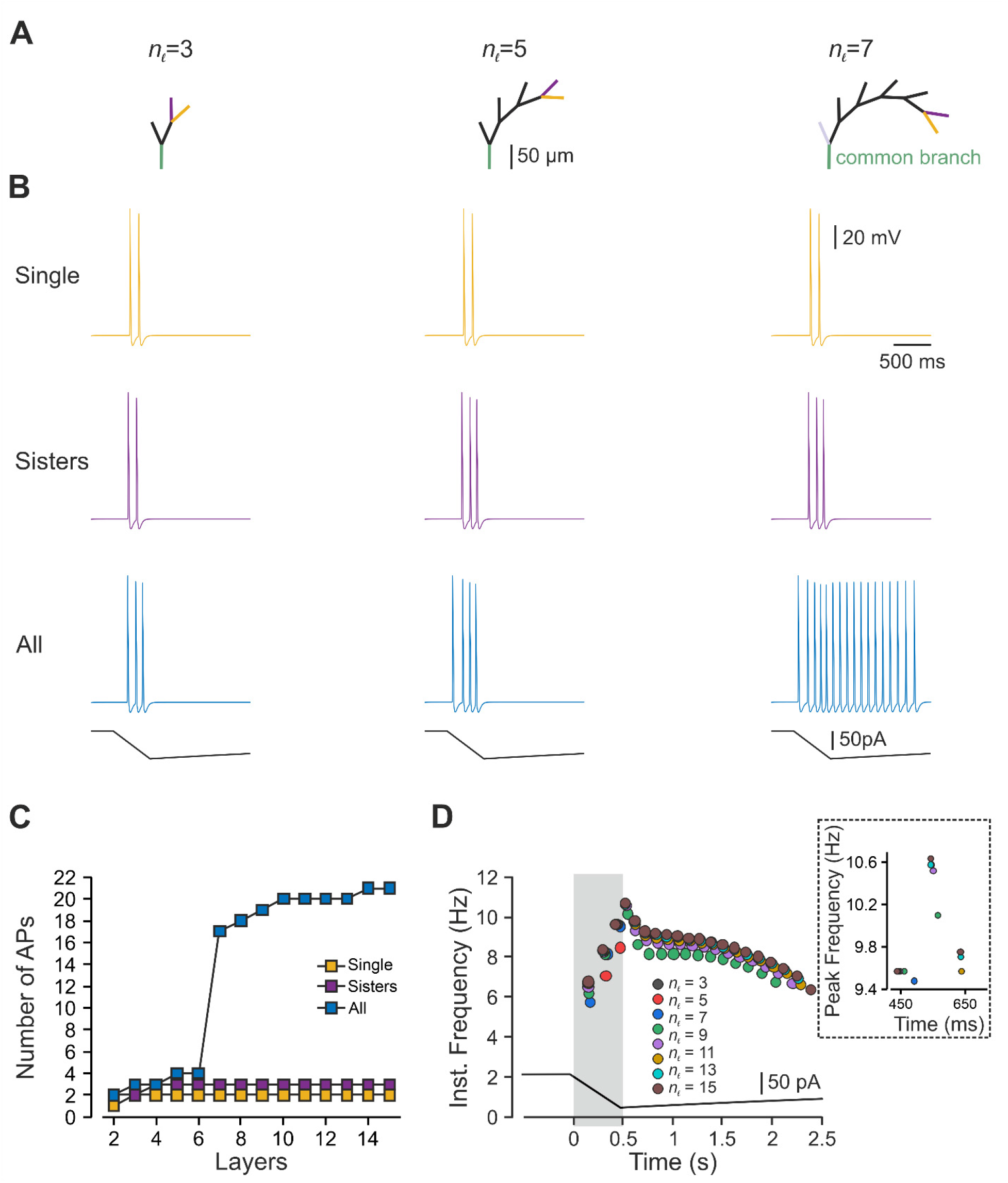
In an asymmetrical degenerate nociceptive terminal tree, an increased number of stimulated terminals facilitates nociceptive response. **A.** Scheme depicting the model of the degenerate binary trees of different complexities determined by the number of bifurcation stages (n*l, see Methods*), which defines the number of terminal branches. **B.** Responses detected at the central terminal following stimulation of either single terminal branch (*yellow*), pair of adjacent terminal branches (*purple*) and all terminals (*blue*) by a capsaicin-like current (depicted below the traces) applied to the representative n_*l*_ = 3, n_*l*_ = 5 and n_*l*_ = 7 trees. **C.** The number of APs plotted as a function of the number of bifurcation stages, following stimulation of a single terminal branch (*yellow*), pair of adjacent terminal branches (*purple*) and all terminals (*blue*) by a capsaicin-like current. **D.** Instantaneous discharge frequencies following stimulation with a capsaicin-like current (*shown below*) applied to all terminals of the trees of different complexity (n_*l*_). Inset: instantaneous peak frequency (color-coded as in *D*).

**Figure 6.**
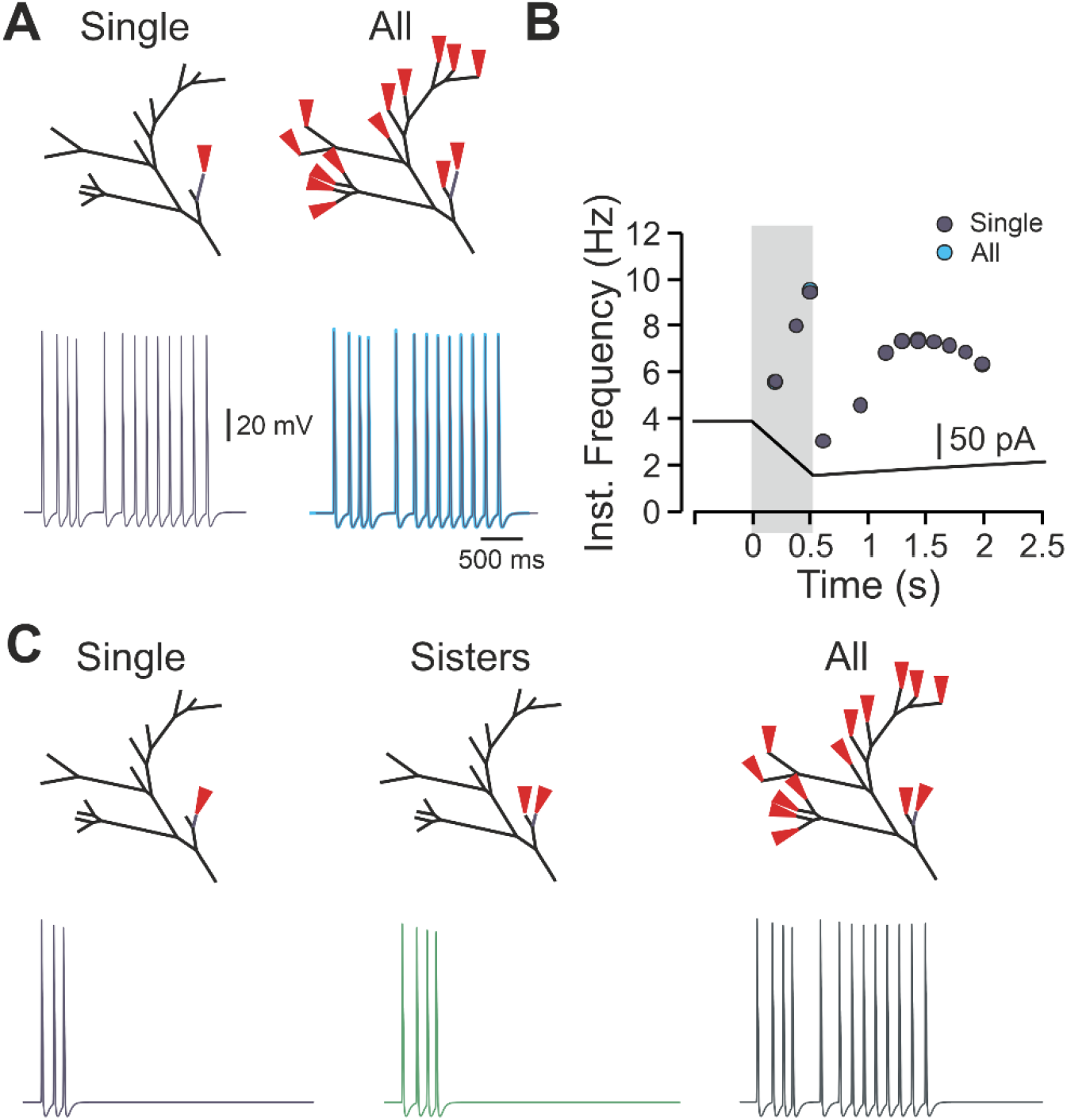
The location of a single terminal branch defines its contribution to the terminal response. **A.** *Upper panels,* schemes depicting the locations of the stimulation. *Left lower panel*, AP firing detected at the central terminal following stimulation of single 175 μm terminal branch located close to the terminal branch (*navy blue terminal at the upper panel*). *Right lower panel*, superimposition of the traces obtained following stimulation of all terminal branches (*light blue*) and stimulation of a single 175 μm terminal branch shown in the left. Note that the firing is identical. **B.** Instantaneous frequencies plotted as a function of time, following stimulation by a capsaicin-like current of a single 175 μm terminal branch shown in A, *left* (*navy blue*), or all terminal branches (*light blue*). Shadowed area outlines the 500 ms of a capsaicin-like current step. **C.** Same as *A,* but the terminal stimulated in *A* was shortened to 75 μm. The stimulation of this terminal branch resulted in the firing of three APs (*lower left*). The stimulation of this terminal branch, together with the similar adjacent terminal branch, led to the generation of four APs (*lower middle*) and was substantially lower than the firing resulted from stimulation of all the terminal branches (*lower right*).

We, therefore, examined the effects of the properties of the individual terminals on the nociceptive gain. Nociceptive terminal trees are composed of terminal branches with different lengths and thicknesses (Zylka et al., 2005; Ivanusic et al., 2013; Alamri et al., 2015; Olson et al., 2017; Alamri et al., 2018; Bouheraoua et al., 2019). The application of a simulated capsaicin current produced a variety of responses depending on the length of the individual stimulated terminals. Stimulation of the shortest single terminal (total length of 75 μm) triggered three APs at the central terminal (Figure 7A). The same stimulation applied to a longer adjacent (distance-wise from the common branch), terminal produced a substantially higher firing at the central terminals (Figure 7B), culminating at 13 APs following stimulation of the longest 175 μm terminal (Figure 7C). The longest terminal was situated at the longest distance from the common branch, suggesting that in addition to the location of the individual terminal, the length of the terminal branch is one of the factors in defining the terminal gain.

**Figure 7.**
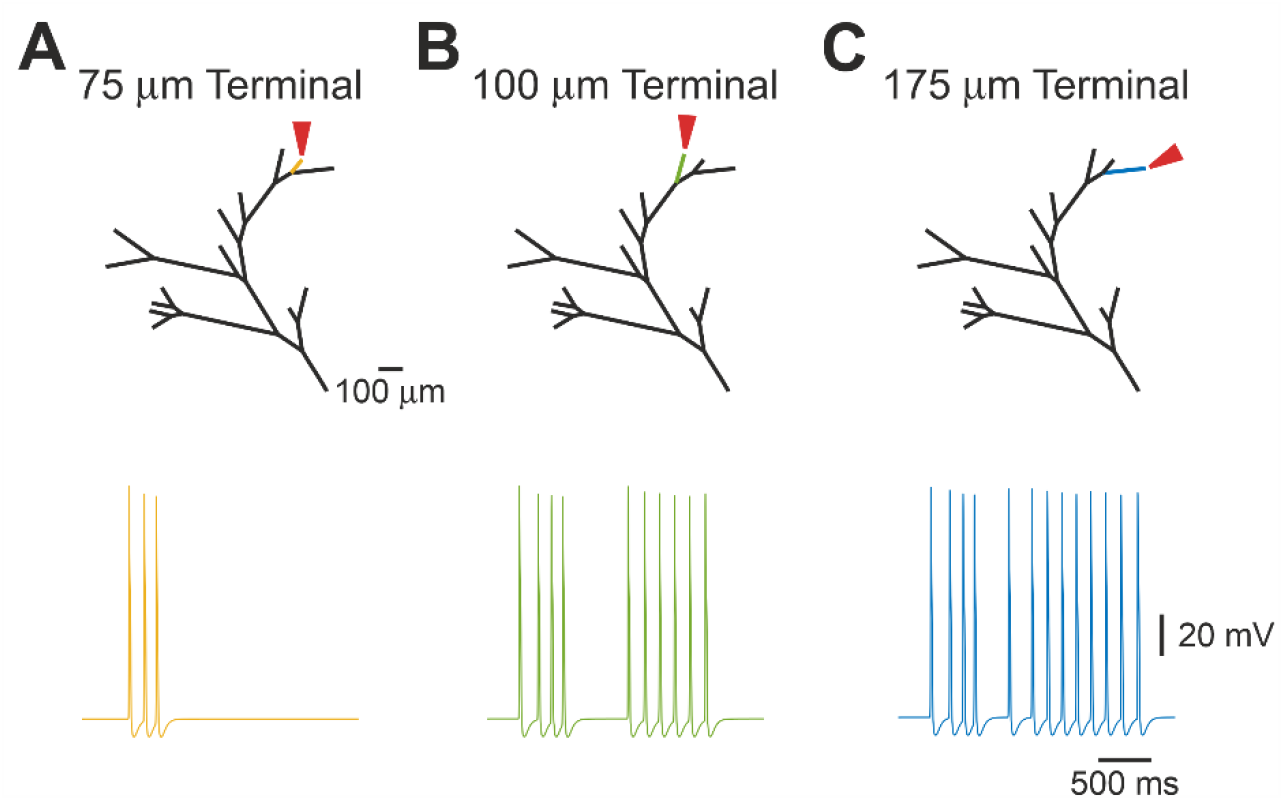
The length of the terminal branch affects the response to a capsaicin-like current. **A-C.** Simulated voltage-clamp recordings from a central terminal following stimulation of single terminal branches of different length (75 μm, *yellow,* **A**, 100 μm, *green,* **B** *, and 1*75 μm, *dark blue,* **C**), by a capsaicin-like current. Note that the firing depends on the lengths of the stimulated terminal branch.

In our model, each terminal consists of two parts: a 25 μm, constant length distal “passive” part, which receives the stimulation, and does not contain Navs (Goldstein et al., 2019). The subsequent “conductive” part contains Navs, and it varies in length before it converges onto the fiber (Figure 1A). In order to reflect the presence of intracellular organelles within the terminal branch (Müller et al., 1996; Bekkers, 2011; Alamri et al., 2018; Goldstein et al., 2019), we increased the axial resistance of the terminal branch (Ra_*TB*_) 15 times that of the value of the fiber (Bekkers, 2011; Goldstein et al., 2019). In this respect, the passive part of the terminals, which receives the stimulus and is connected to the highly resistive conductive part, is somehow similar to a “head” of the dendritic spine connected to the spine neck (Nimchinsky et al., 2002). Accordingly, the voltage response of the passive part to a capsaicin-like current is dependent on its own resistive properties, which define the resulting voltage deflection, according to Ohm’s law. However, it may also depend on the axial resistance (R_a_) of the consecutive conductive part, such that, similarly to a spine head and neck, a high Ra component of the conductive part will lead to higher voltage deflection at the SIZ and hence a higher firing, by effectively decreasing the current sink (Segev and Rall, 1988). Consequently, we examined capsaicin-induced voltage deflection along the terminal branch, by stimulating the tip of the terminal by a capsaicin-like current and monitoring the resulting voltage deflection at the “passive” part, 2.5 μm before the SIZ (Figure 8A) and at the SIZ, while changing the Ra of the conductive part (Figure 8A, B). We removed all Nav conductances from the conductive part, to eliminate the effect of Nav activation and APs on the voltage deflection. To examine how voltage deflection at the SIZ affects the AP firing at the central terminal we added back on the Nav conductances (Figure 8C). When the conductive part Ra value was 15 times that of the fiber, stimulation of the terminal tip with a capsaicin-like current produced voltage deflection of about 60 mV, depolarizing the membrane potential (Vm) of the terminal end to 0 mV which is equivalent to the TRPV1’s reversal potential (Figure 8A). At the SIZ the voltage deflection was smaller, probably due to the voltage decay resulting from a high space constant, reaching VmSIZ of –12 mV (Figure 8B). This depolarization at the SIZ was sufficient to produce an excessive firing at the central terminal (13 APs, Figure 8C). Reduction of the conductive part’s Ra to one times of the fiber led to a smaller depolarization (ΔVm_*Tip*_ of 35 mV) at the terminal tip (Figure 8A) and at the SIZ (ΔVm_*SIZ*_ of 25 mV, Figure 8B). At these conditions no APs were evoked by stimulation with capsaicin-like current when Nav conductances were turned back on (Figure 8C). These data suggest the changes in Ra solely in the conductive part, strongly affect the terminal membrane depolarization and AP firing.

**Figure 8.**
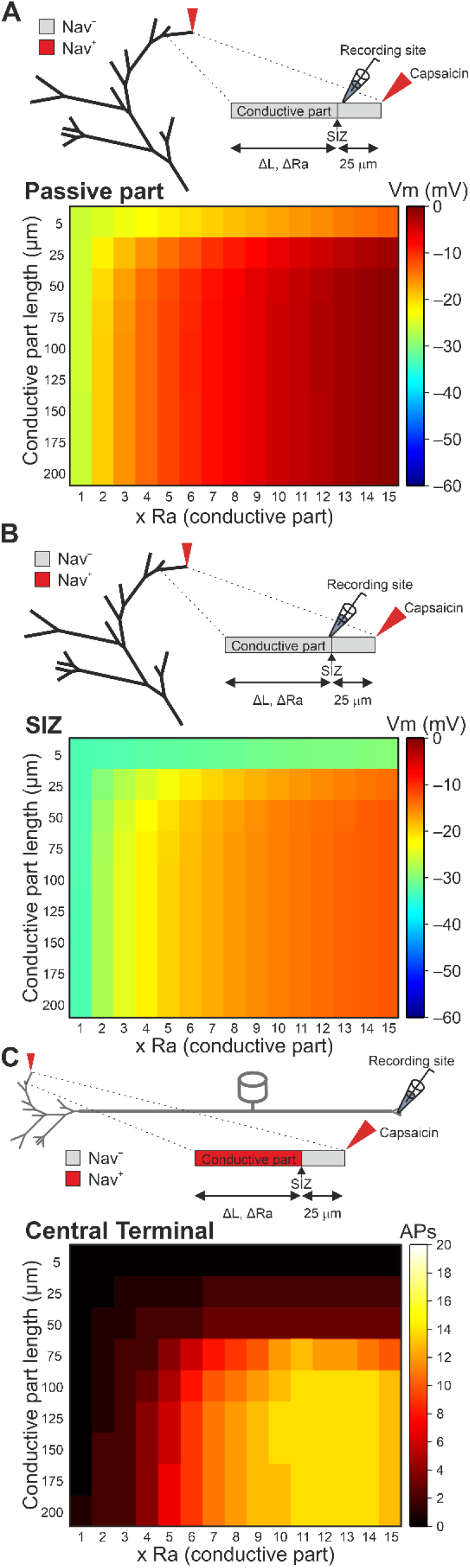
The axial resistance of the terminal branch’s conductive part determines the nociceptive gain. **A.** *Upper*, the scheme of the stimulated terminal branch with the location of the recording pipette 2.5 μm before the SIZ. The Nav conductances at the conductive part were annulled to prevent the effect of the APs on the measurements of the terminal membrane potential. *Lower*, the heatmap representing the relation between the axial resistance of the conductive part (Ra), the conductive part length, and the depolarization of the terminal membrane recorded at the Nav-less part, 2.5 μm before the SIZ. The resulted membrane potential is color-coded (shown on the *right*). Note that when the Ra of the conductive part is similar to the Ra of the proximal fiber (1 X Ra), the depolarization resulted from the application of a capsaicin-like current to the terminal tip is substantially lower than when Ra is higher. **B.** Same as *A*, but the recordings are performed at the SIZ. **C.** The relation between the axial resistance of the conductive part (Ra), the conductive part length, and the AP firing measured at the central terminal following stimulation of a single terminal. The Nav conductance of the conductive part is intact. The AP count is color-coded (shown on the *right*).

These results, however, do not explain the increase in firing when longer terminals are stimulated (Figure 7) since all the terminals had a similar Ra. Indeed, the effect of length on the firing at the central terminals was independent of Ra values, and at any given Ra the change in the length of the conductive part affected the firing at the central terminal in a sigmoidal manner as the length increased (Figure 8C). The shortening of the “conductive” terminal part below 25 μm prevented the generation of APs at all Ra values, whereas the elongation of the conductive part leads to the increase in the firing at the central terminal which peaked and plateaued, when the conductive part was of 125-150 μm (Figure 8C). It has been shown that the length of the SIZ of central neurons defines neuronal excitability, by virtue of the amount of Nav conductance (Kuba et al., 2010). Our model suggests that an increase in the length of the Nav containing components enhances nociceptive gain (Figure 4). Consequently, we examined whether the change in length of the terminals regulates the terminal gain by affecting the Nav conductance. Nociceptive neurons express a variety of sodium channels. Among them, slow Nav 1.8-mediated sodium current defines the ability of a nociceptive neuron to fire repetitively (Blair and Bean, 2003). Persistent Nav 1.9-mediated sodium current controls subthreshold excitability (Cummins et al., 1999; Herzog et al., 2001; Baker et al., 2003). Both Nav 1.8 and 1.9 are expressed by nociceptive terminals (Persson et al., 2010). We stimulated a fixed-length terminal by a capsaicin-like current and mimicked the change of the terminal length by varying either Nav1.8 or Nav1.9 conductances. Increase in Nav 1.8 conductance to 145% in short terminals of 50 μm (25 μm Nav-less compartment + 25 μm conductive, Nav expressing, compartment), which generates two APs in the control conditions, lead to the firing of 13 APs, which is comparable with the firing generated by a 125 μm long terminal (Figure 9A). In a 75 μm long terminal (25 μm Nav-less compartment + 50 μm conductive, Nav expressing, compartment), which fires 3 APs in normal conditions, the increase of Nav1.8 conductance to 110% was sufficient to generate 14 APs, which is comparable with the firing of a 125 μm terminal (Figure 9A). Further increase in Nav1.8 conductance leads to higher firing, exceeding that of longer terminals with “normal” Nav1.8 conductance (Figure 9B). Shortening of the terminal, by decreasing Nav1.8 conductance, reduces the firing in an abrupt manner, such that an application of capsaicin-like current to the terminal with 95% of Nav1.8 conductance leads to the generation of 4 instead of 13 APs and a decrease of Nav1.8 conductance below 55% prevented AP generation (Figure 9B). Changing only in Nav 1.9 conductance resulted in a more gradual effect on AP firing (Figure 9C, D), and in short terminals of 50 and 75 μm, the increase in Nav1.9 conductance to 150% was not sufficient to increase the terminal gain properties (Figure 9D). In a longer terminal of 100 μm, an increase in Nav1.9 conductance by 10% increased the firing to the levels similar to the firing of longer terminals (Figure 9C). Similar to the Nav1.8 conductance, a further increase in Nav1.9 conductance in terminals longer than 100 μm produces substantially higher firing than the firing of the long terminals in “normal” conditions (Figure 9D). The decrease in Nav 1.9 conductance led to a gradual reduction of the firing, which was reduced but not annulled even when Nav1.9 conductance was zeroed (Figure 9D). These data suggest that longer terminals are more excitable due to higher expression of Nav1.8 and Nav 1.9 channels. The Nav 1.8 conductance defines the repetitive firing, and Nav1.9 tunes it. The non-linearity between Nav1.8 and 1.9 conductances, the length, and the firing, could be explained by the effect of other conductances regulating the firing. Since we only manipulated Nav conductance and not the length of the conductive part, our manipulation left all other conductances intact; thus, an increase in Nav1.8 or Nav1.9 conductance was not counteracted by the increased potassium conductances, which normally restrain the firing.

**Figure 9.**
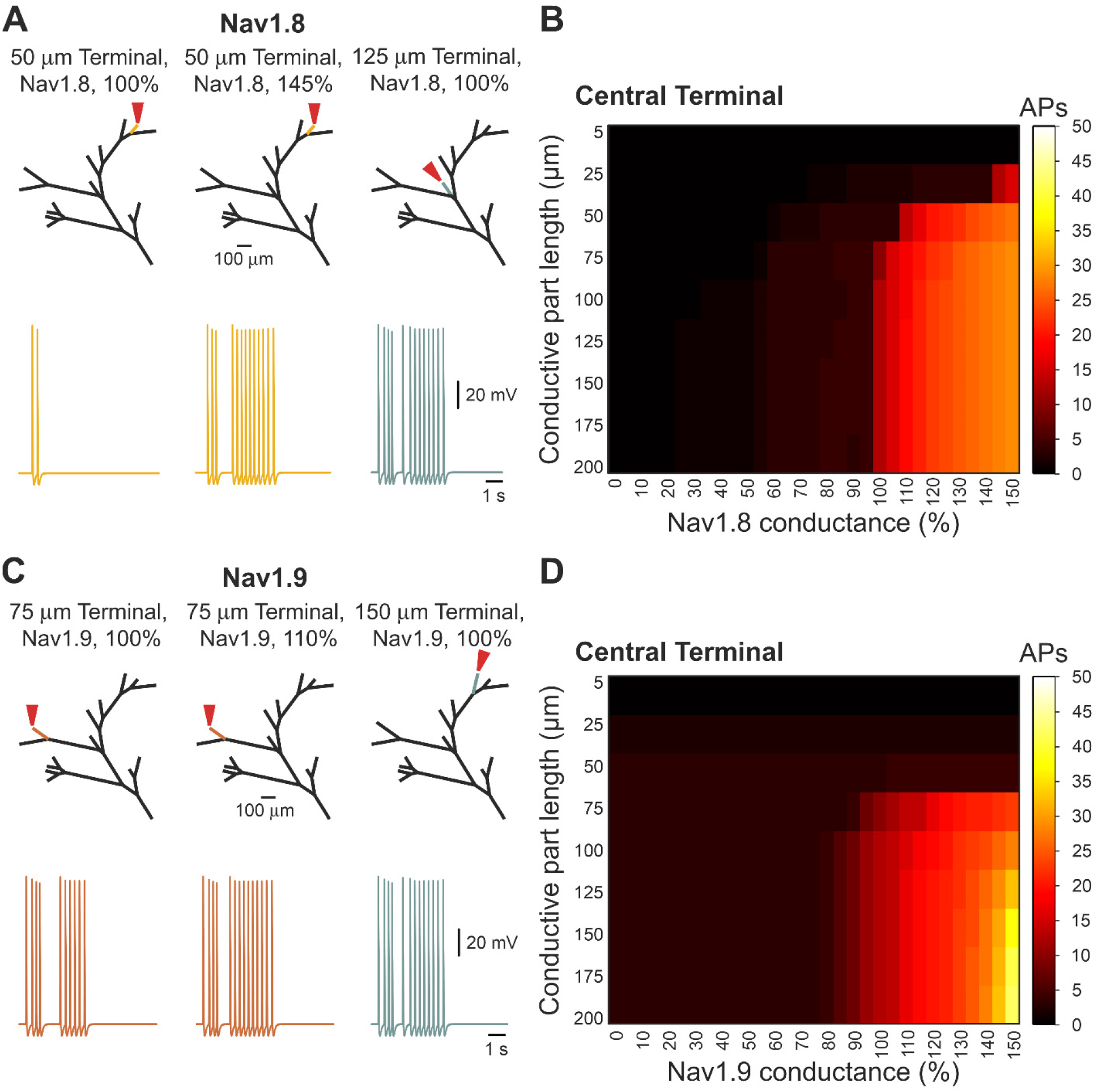
The length of the conductive parts affects the nociceptive gain by defining the level of Nav 1.8 and Nav 1.9 conductances. **A.** *Left*, a trace of the response to the stimulation (*red arrow*) of the 50 μm terminal branch (*yellow*) with the Nav1.8 conductance at the control level (100%). Stimulation of the same terminal branch but with increased Nav1.8 (145%) conductance led to higher firing (*middle*), which was compatible with the stimulation of longer (125 μm) terminal branch with the control (100%) Nav1.8 conductance (*right*). **B.** The heatmap representing the relation between the length of the conductive part, the relative level of Nav conductance, and the AP firing measured at the central terminal following stimulation of a single terminal. The AP count is color-coded (shown on the *right*). Note that firing following the stimulation of the terminals with the 100% Nav 1.8 conductance is increased when the Nav conductance is increased. Note also that the decrease in Nav 1.8 conductance in the long terminal branches, which normally triggers a high number of APs, led to a dramatic reduction of firing. **C and D.** Same as A and B but showing the relation between Nav1.9 conductance, the conductive part length, and the AP firing at the central terminal.

Altogether these data suggest that the properties of the individual terminals and the complexity of the terminal tree affect the gain of nociceptive input-output function. The value of the generation potential is defined by the resistance of the terminal branch. The translation of the generation potential into the AP firing depends on the length of the Nav 1.8 and 1.9 expressing compartments and the integration of firing from multiple branches. All these factors define the encoding properties of the nociceptive terminal.

In all the experiments described above, we stimulated all the branches simultaneously. However, in physiological conditions, the stimulus (e.g., heat, cold, capsaicin) applied to a certain region area dissipates from the center of the stimulated areas into the surroundings; thus, each terminal receives the stimuli at the different time, possibly leading to a temporal summation of the stimuli. We, therefore, studied whether the temporal integration of noxious stimuli amplifies the nociceptor excitability. To simulate a stimulation of the receptive field with respect to time variance among terminals, we reformulated a capsaicin-like stimulations with the addition of a normally distributed probability randomized onset time variability (Figure 10A). The simulations were then applied to all terminals of a complex n_*l*_ = 7 balanced binary type terminal tree (Figure 10B). An increase in onset time probability from 10 to 2500 ms resulted in an exponential increase of total AP firing recorded in the central terminal over logarithmically increased onset time variance (Figure 10C). Changes between 10 to 250 ms produced only a small increase in total AP firing over time (<20). However, the extension of onset time probability greatly increased the excitatory output (Figure 10C) up to a 2-fold increase in longer onset times (Figure 10C). These data indicate that the prolonged continuous capsaicin-like currents stimuli feature a temporal-summation characteristic such that the successive application of these stimuli leads to an increase in the response of nociceptive neurons. Our data also implies that the facilitation we observed in the degenerate binary tree (Figure 5) could be due to the temporal integration of the responses from different terminals.

**Figure 10.**
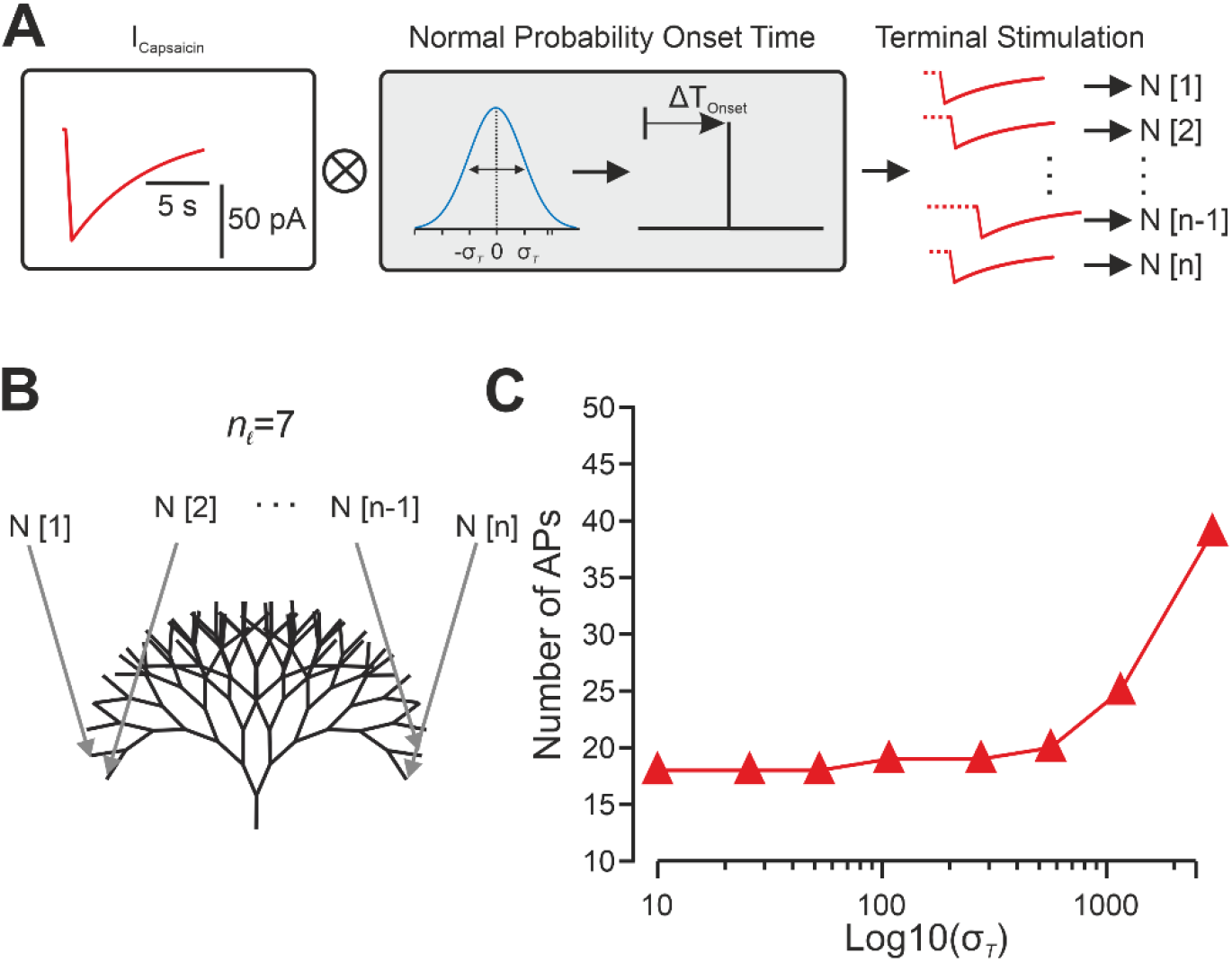
Temporal summation by the nociceptive terminal tree. **A.** Scheme of normally distributed random time onset for a single of *n* terminal stimulations (*N*). After a capsaicin current is generated (*red*) its stimulation onset time is shifted to a normal distribution (*blue*) random onset time with a standard deviation (σ). *n* stimulations are then each applied to a single terminal index with respect to its index (*1‥n*). **B.** Scheme depicting the model of a 7 bifurcation stages balanced binary tree. Each of the time-randomized stimulations (*N[1‥n]*) is applied to a terminal nerve-ending (*grey arrows*). **C.** The number of APs generated as a function of the logarithmic scaling of normal distribution standard deviation onset time of a capsaicin-like current in all terminals. The activity was measured at the central terminal.

While the simulations described above were done in “normal” conditions, we next examined how nociceptive terminals integrate the increased responses to noxious stimuli in pathological conditions. Many pathological conditions lead to hyperalgesia, i.e‥, increased pain response to noxious stimuli (Sandkuhler, 2009). We previously demonstrated that inflammation leads to hyperalgesia by shifting the location of the SIZ of the nociceptive terminals closer to the terminal tip (Goldstein et al., 2019). Consequently, we shifted the location of the SIZ towards the terminal tip (Figure 11A), to simulate the changes which occur during inflammatory hyperalgesia. Activation of all terminals with a capsaicin-like current under these conditions slightly increased the firing (15 instead of 13 APs, Figure 11B) and the peak instantaneous frequency from 9.4 Hz to 10.4 Hz (Figure 11C). To simulated excitability changes in neuropathic pain, we conferred upon nociceptive membrane sub-and suprathreshold perturbations of the membrane potential (Barkai et al., 2017). It has been demonstrated that nerve injury leads to an increase in the resting membrane potential voltage perturbations, which may initiate spontaneous ectopic activity (Amir et al., 1999; Liu et al., 2000; Rho and Prescott, 2012) and that this ectopic activity is correlated with neuropathic pain (Kleggetveit et al., 2012). Accordingly, we have injected an Onstein-Uhlenbeck based current noise (Olivares et al., 2015) in which the amplitude of the injected current varies according to the normal distribution at each time step. We simulated subthreshold noise and suprathreshold spontaneous activity by varying the standard deviation of the current amplitude’s normal distribution (σ) between 0.006 to 0.007, respectively (Figure 11A). In the higher σ of 0.007, the probability of reaching higher injected currents was higher; thus, it caused higher membrane fluctuation, which sometimes reached the threshold, and generated AP (Figure 11A). In these conditions, activation of all terminals by a capsaicin-like current substantially increased firing (Figure 11B) and the peak instantaneous frequency was slightly increased (9.9 Hz in subthreshold noise and 10.1 Hz in suprathreshold noise, Figure 11C).

**Figure 11.**
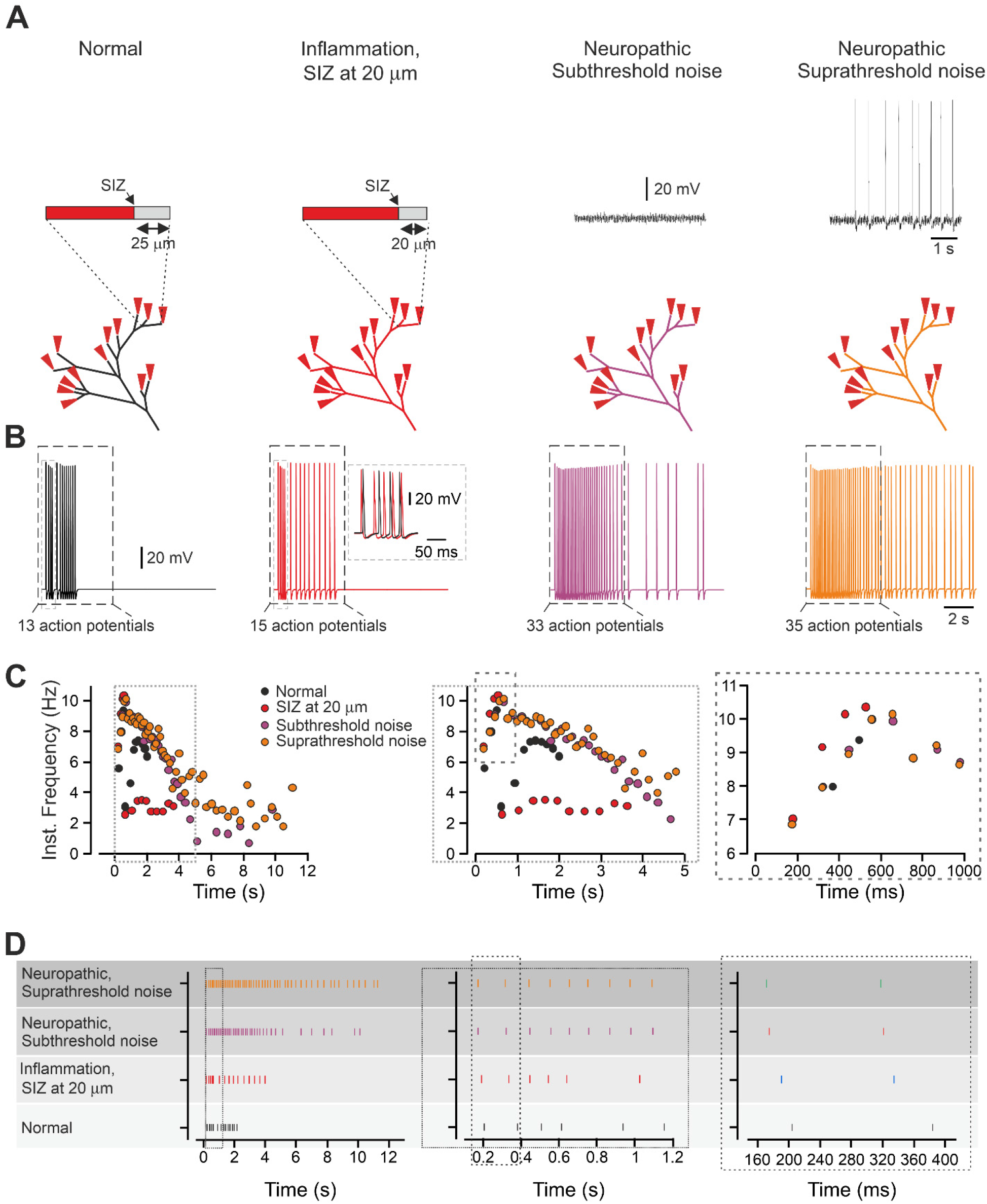
In modeled inflammatory and neuropathic conditions, the change in response is encoded by alteration in the temporal pattern of the spike trains. **A.** Scheme of the experiment: in the “normal” condition, the SIZ is located 25 μm from the terminal end, and the terminal membrane potential is quiescent. The inflammatory hyperalgesia is modeled by shifting the SIZ 5 μm towards the terminal tip. The neuropathic hyperalgesia is modeled by injected to the model sub-or suprathreshold current noise (*see Methods*). The insets show the recording of the membrane voltage following the application of sub-and suprathreshold noise. **B.** Responses detected at the central terminals following activation of all terminal branches by a capsaicin-like current in normal conditions (*black*), in the conditions of the inflammatory hyperalgesia (*red*) and in the conditions of the neuropathic hyperalgesia with the injection of subthreshold noise (*purple*) and suprathreshold noise (*orange*). The AP count was performed for a first 5 s after a capsaicin-like current application (outlined by the dashed-line rectangle). *Inset*: superimposition of the traces obtained in normal conditions and in the inflammatory hyperalgesia conditions at the time outlined by the gray dashed-line rectangle. Note the difference in the pattern of firing. **C.** Instantaneous discharge frequencies following stimulation with a capsaicin-like current (*shown below*) applied to all terminal branches in normal conditions (*black*), in the conditions of the inflammatory hyperalgesia (*red*) and in the conditions of the neuropathic hyperalgesia with the injection of subthreshold noise (*purple*) and suprathreshold noise (*orange*). *Insets* show expanded x-axis. **D.** A spike raster plot showing the AP timing, following stimulation of all terminal branches with a capsaicin-like current, in normal conditions (*black*), in the conditions of the inflammatory hyperalgesia (*red*) and in the conditions of the neuropathic hyperalgesia with the injection of subthreshold noise (*purple*) and suprathreshold noise (*orange*). Insets show expanded x-axis. The data describing the normal conditions are taken from Figure 1 and showed here for better comparison.

Our data demonstrate that activation of all terminals of the “realistic” terminal tree in normal conditions led to the generation of 13 APs, which fires at the central terminal with an instantaneous frequency peaked at 9.3 Hz (Figure 1E and 11C). Activation of complex balanced or degenerate trees, although produced a higher firing (18 and 21 APs, respectively), the peak instantaneous frequency remained within a similar limit of 9 to 10 Hz (Figures 3 and 5). Similarly, in the simulated pathological conditions, the peak instantaneous frequency was only slightly increased (*see above*). This change in peak instantaneous frequency may be too narrow to encode and convey the changes in response to stimuli. We, therefore, hypothesized that in addition to the spike rate coding (Adrian and Zotterman, 1926), the nociceptive neurons might also use “temporal” coding to integrate and encode the information about different response properties via different timing patterns of spikes (Cho et al., 2016). Accordingly, we analyzed the spiking activity over time following a capsaicin-current stimulation of all terminals in the “realistic” terminal tree in normal conditions, in inflammatory hyperalgesia conditions, in which the SIZ was shifted toward the terminals and in neuropathic pain conditions, in which sub- and suprathreshold current noise was injected (Figure 11D). The simulation revealed that the peak discharge rate following the stimulation at different conditions was almost unchanged, while the patterns of the discharges evoked at different conditions were substantially different (Figure 11D). These data suggest that changes in temporal encoding of the spike trains from an individual nociceptive neuron may convey key information about changes in nociceptive output in pathological conditions, thus defining abnormal nociceptive behavior after nerve injury or inflammation.

In conclusion, our model predicts that the morphology of the terminal tree affects the response of the nociceptive terminal to noxious stimuli. To summarize the effects of the different normal and abnormal morphology of nociceptive terminal tree on the nociceptive functions, we produced a simplified model of “realistic” nociceptive terminal tree which contain “degenerative” and “symmetric” component steaming from the common branch in which all terminal branches are of the same length (75 μm, Figure 12A, *black*). Under these “normal” conditions Activation of this terminal tree by a capsaicin-like stimulus, i.e., simultaneous activation of all terminal branches by the capsaicin-like current generated 10 APs. We then simulated the morphological changes in the terminal branches and the terminal tree and examined how these changes affect the terminal output. An increase in the number of terminal branches to 16, simulating hyperinnervation changes (Cain et al., 2001; Leibovich et al., 2020) leads to an increase in firing (Figure 12A, *red*). A decrease in the number of the terminal branches to 7, simulating a denervation state, substantially decreases the nociceptive gain (Figure 12A, *blue*). Changes in the properties of individual terminal branches by shortening and decreasing Ra of the conductive part also substantially decrease nociceptive gain (Figure 12A, *green*). An increase in the terminal branch length has the opposite effect on the nociceptive gain. Importantly, all the changes in the morphology of the nociceptive terminal tree affected also the pattern of firing (Figure 12B, C, *green*), suggesting that in addition to the changes in the number of the APs, there is also a change in their timing for reaching the central terminal, which is an important factor in nociceptive neuronal processing.

**Figure 12.**
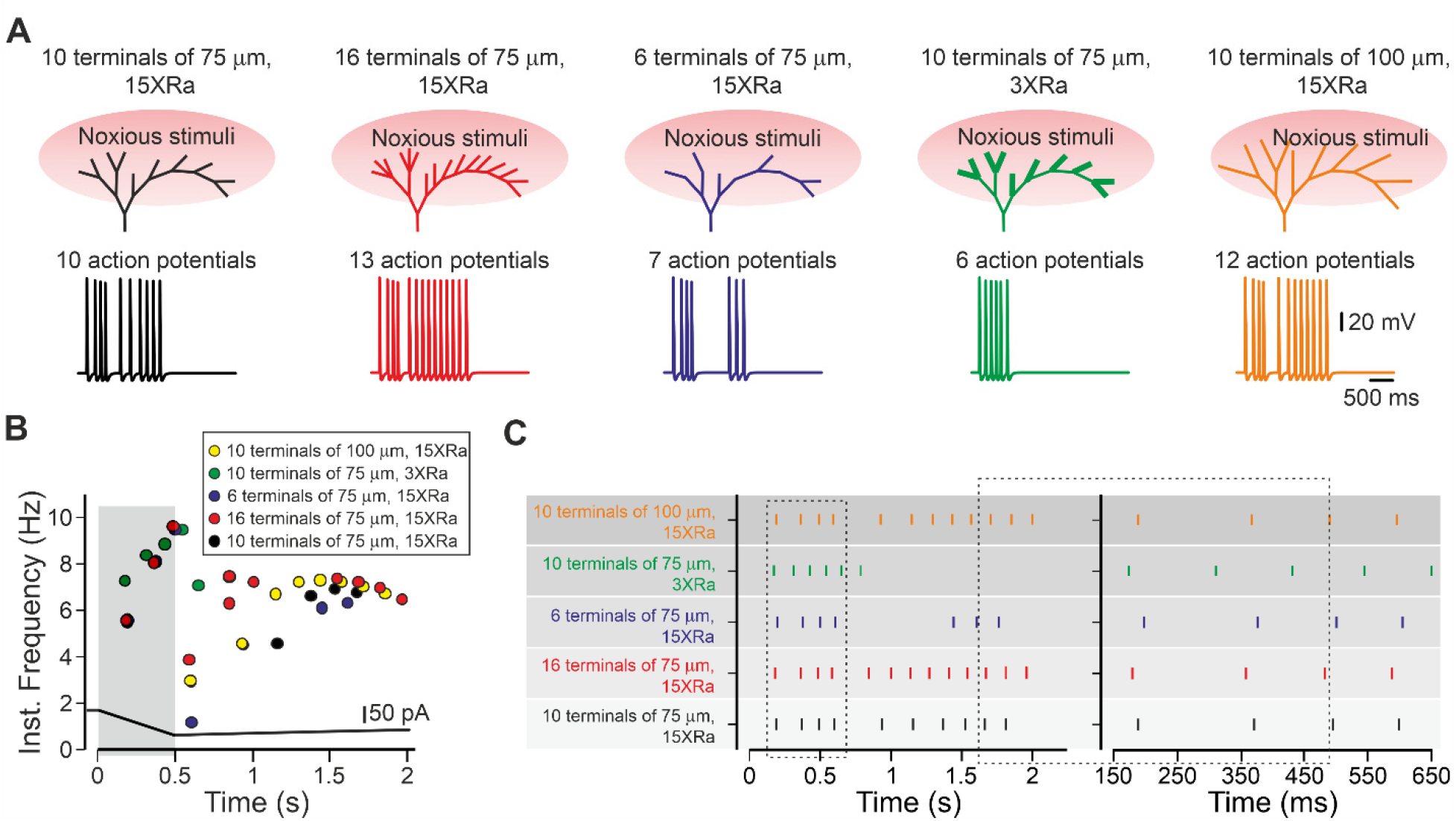
The effect of the morphology of the nociceptive terminal tree on the input-output properties of the primary nociceptive neuron. A figure summarizing the effect of the terminal structure on its function. Activation of all terminal branches in a simplified “realistic” terminal tree (all the terminal branches are of equal size of 75 μm, and they are evenly distributed along the degenerate and symmetrical parts of the complex terminal, *black*) by capsaicin-like stimulus led to the response whose gain (**A**, *lower panels*) and AP temporal pattern (**B, C,** showing the instantaneous frequency and spike raster plot, respectively) are depending on the terminal morphological characteristics, such as change on the number in terminal branches and the properties of the individual branch. The increase in the number of activated terminal branches (*red*) led to increased firing; the decrease in branching (*blue*) led to decreased firing. Terminal tree with short and thick (lower conductive part’s Ra) terminal branches demonstrate lower gain (*green*), whereas a terminal tree with long terminals has a higher gain (*orange*). All the perturbations in terminal morphology change the timing pattern of firing (**B, C**).

## Discussion

Nociceptive terminals innervate the tissue forming a variety of morphologically complex structures (Zylka et al., 2005; Ivanusic et al., 2013; Alamri et al., 2015; Olson et al., 2017; Alamri et al., 2018; Bouheraoua et al., 2019). To examine the effect of the architecture of the whole terminal tree, together with morphological and biophysical properties of its specific component on the nociceptive input-output relation, we used a computational model. This model allowed us to make predictions based on integrating experimental data from a single terminal, fiber, and cell body. We dynamically changed specific parameters of an individual terminal and terminal tree morphology and tested their overall influence on nociceptive input-output relation. Our data predict that the morphology of the terminal tree affects the response of the nociceptive terminal and, thus, the whole nociceptive neuron to noxious stimuli.

We show that the properties of an individual terminal have a prominent effect on nociception and that the single terminal gain is greatly determined by at least two factors. The axial resistance determines the voltage response to the activating current. The resulting AP firing is defined by the length of the conductive part of the terminal, via the increase in the amount of the Navs. These features rendering the nociceptive terminal as a special unit combining properties of two structures known to affect the neuronal gain: a dendritic spine, which increases its synaptic gain by increasing spine-neck axial resistance (Segev and Rall, 1988; Beaulieu-Laroche and Harnett, 2018), and the AIS, a conductive compartment showed to elevate neural excitability with its elongation (Kuba et al., 2010). It is worth mentioning that while these two structures co-exist in a variety of neurons, they are functionally and topologically distant from each other, with spines defining the dendritic input function and the AIS attributed to axonal output function. Here, we show that the nociceptive terminal is a bi-functional compartment combining these two functional characteristics into one compartment, the terminal branch, outlining its importance in defining the nociceptive gain. Interestingly, both spine-neck Ra and AIS length are considered key to neuronal plasticity (Magee, 2000; Grunditz et al., 2008; Sjöström et al., 2008; Grubb et al., 2011; Araya et al., 2014; Gulledge and Bravo, 2016; Yamada and Kuba, 2016). It is, therefore, reasonable to consider terminals as key regulatory sites for nociceptive plasticity. Thus, changes in the electrical properties of a terminal (Goldstein et al., 2019) could lead to higher or lower sensitivity to pain. For example, our data predict that the change in the density of intracellular organelles at the terminal branch can cause a local increase of the terminal axial resistance and thereby an increase in its excitatory characteristics, leading to increased pain. The changes in the electrical properties of a single terminal leading to an increase of its gain could serve as a possible explanation to the state in which a decrease in fiber density due to nerve degeneration leads to increased pain (Chiang et al., 2018). Previous studies on central neurons demonstrated the relation between axonal morphology and neuronal activity (Manor et al., 1991; Ofer et al., 2017). These studies examined the effect of the axonal structure on the efferent propagation of the action potentials. Here we examined the integration of afferently propagating signals and how they merge at the convergence points. Moreover, the majority of the studies examined the effect of short stimuli applied at different frequencies. Our data predict no summation of the afferent information within the terminal trees when short stimuli are applied (Figure 1). However, we show a substantial voltage summation at the convergence points when more “natural” capsaicin-like prolonged stimulation is applied, suggesting that “natural” activation of terminal trees with more terminal branches might lead to increased nociceptive output and thus to increased pain.

It was recently demonstrated that neuropathic pain following nerve injury was associated with a decreased number of DRG neuronal somata innervating the hypersensitive area (Leibovich et al., 2020) and increased number of axons innervating the area (Duraku et al., 2012). Together these results suggest that each DRG neuron innervating the hypersensitive area increases the complexity of its terminal fibers. Our model predicts that the increase in the terminal branching of individual DRG neurons following nerve injury (Leibovich et al., 2020) or inflammation or tumor formation (Cain et al., 2001) affects neuronal excitability and thus may contribute to the development of pain hypersensitivity in neuropathic conditions.

We show that in addition to a spatial summation at the convergence points, the physiological activation of nociceptive terminals also permits temporal summation of the stimuli. This temporal summation is unveiled due to the asynchronous activation of the terminals in the physiological conditions. In the case of nociceptive neurons with large receptive fields such as of the back and leg (Mancini et al., 2014), the stimuli will first activate the terminals at the epicenter of the stimuli and only then at the surrounding areas as it dissipates. But it is also true in the case of nociceptors with small receptive fields of palms and fingers (Mancini et al., 2014), as terminals vary in skin depth and distance from the stimulation point (Dezhdar et al., 2015; Alamri et al., 2018). Therefore, a single terminal stimulation onset time differs between the terminals covering either large and also small areas. At the dendrites, the differences in location and time of stimulation define the resulting output of the stimulated neuron (Magee, 2000; Magee and Johnston, 2005). Traditionally, the temporal proximity of a series of subthreshold synaptic events can, as a result of EPSP temporal integration, increase the output of the postsynaptic cell in terms of its firing probability and AP frequency. We show that activation of complex nociceptive terminals allows temporal summation, leading to response amplification. Importantly, this summation, unlike summation of postsynaptic potentials, is not strictly a passive property of the membrane but rather results from an active post-AP prolonged subthreshold depolarization, which allows postsynaptic potential-like “passive” summation.

Neurons encode information using both the spike rate coding (Adrian and Zotterman, 1926) and the temporal coding (Panzeri et al., 2001; Johansson and Birznieks, 2004). Our model predicts that in nociceptive neurons, different excitability conditions resulting in a slight difference in peak instantaneous frequency. These differences in spike rate may be too narrow to encode the differences in neuronal excitability. We show that in modeled pathological conditions in which nociceptive neurons become hyperexcitable, the changes in their gain are encoded by the differences in the spike arrival time. The differences in spike patterns between capsaicin and GABA-evoked responses were recently correlated with the differences in the resulting pain sensation following either capsaicin or GABA stimulation (Cho et al., 2016). Thus, the difference in the timing pattern of spikes could convey important information regarding the gain of the input-output properties of the nociceptive neurons.

In summary, we introduced here a realistic model of the nociceptive terminal tree, which reveals that the structure of the terminal tree defines the function of nociceptive neurons. As in all models, although it does not reflect all neuronal morphological and physiological properties, its simplicity allows examining how a change in a specific parameter affects the whole complex multiparameter system. Accordingly, our model permits the precise, repeated, time-locked stimulation of a designated neuronal compartment of the nociceptive neuron. It also could be utilized to test other hypotheses of the physiology and pathophysiology of nociception. For example, we used a capsaicin-like stimulus; however, our model could be adapted to predict how nociceptive neurons respond to other types of noxious stimuli and how these responses vary depending on changes in different ion conductances, electrical properties of the terminal, terminal architecture or under pathological conditions. Our model predicts how activation of the nociceptive neurons with different terminal architecture would change the neuronal response. It also suggests that the changes in the tree architecture in pathological conditions may be sufficient to modify the input-output relation of the primary nociceptive neurons, thus leading to pathological pain. The knowledge collected from our model gives rise to predictions and speculations, which otherwise could not be achieved with the current experimental tools. Moreover, the results of this theoretical study open new avenues to follow and explore experimentally.

## Acknowledgments

Support is gratefully acknowledged from the Israeli Science Foundation - grant agreement 1470/17; Canadian Institutes of Health Research (CIHR), the International Development Research Centre (IDRC), the Israel Science Foundation (ISF) and the Azrieli Foundation - grant agreement 2545/18; the Deutsch-Israelische Projectkooperation program of the Deutsche Forschungsgemeinschaft (DIP) grant agreement BI 1665/1-1ZI1172/12-1

## References

Adrian ED, Zotterman Y (1926) The impulses produced by sensory nerve-endings: Part II. The response of a Single End-Organ. J Physiol 61:151–171.

Alamri A, Bron R, Brock JA, Ivanusic JJ (2015) Transient receptor potential cation channel subfamily V member 1 expressing corneal sensory neurons can be subdivided into at least three subpopulations. Frontiers in Neuroanatomy 9.

Alamri AS, Wood RJ, Ivanusic JJ, Brock JA (2018) The neurochemistry and morphology of functionally identified corneal polymodal nociceptors and cold thermoreceptors. PLoS One 13.

Amir R, Michaelis M, Devor M (1999) Membrane potential oscillations in dorsal root ganglion neurons: role in normal electrogenesis and neuropathic pain. J Neurosci 19:8589–8596.

Araya R, Vogels TP, Yuste R (2014) Activity-dependent dendritic spine neck changes are correlated with synaptic strength. Proc Natl Acad Sci U S A 111:30.

Baker MD (2005) Protein kinase C mediates up-regulation of tetrodotoxin-resistant, persistent Na+ current in rat and mouse sensory neurones. J Physiol 567:851–867.

Baker MD, Chandra SY, Ding Y, Waxman SG, Wood JN (2003) GTP-induced tetrodotoxin-resistant Na+ current regulates excitability in mouse and rat small diameter sensory neurones. J Physiol 548:373–382.

Barkai O, Goldstein RH, Caspi Y, Katz B, Lev S, Binshtok AM (2017) The Role of Kv7/M Potassium Channels in Controlling Ectopic Firing in Nociceptors. Front Mol Neurosci 10:181.

Basbaum AI, Bautista DM, Scherrer G, Julius D (2009) Cellular and molecular mechanisms of pain. Cell 139:267–284.

Beaulieu-Laroche L, Harnett MT (2018) Dendritic Spines Prevent Synaptic Voltage Clamp. Neuron 97:75–82.

Bekkers JM (2011) Changes in dendritic axial resistance alter synaptic integration in cerebellar Purkinje cells. Biophys J 100:1198–1206.

Belmonte C, Aracil A, Acosta MC, Luna C, Gallar J (2004) Nerves and sensations from the eye surface. Ocul Surf 2:248–253.

Binshtok AM (2011) Mechanisms of nociceptive transduction and transmission: a machinery for pain sensation and tools for selective analgesia. International review of neurobiology 97:143–177.

Blair NT, Bean BP (2002) Roles of Tetrodotoxin (TTX)-Sensitive Na Current, TTX-Resistant Na Current, and Ca Current in the Action Potentials of Nociceptive Sensory Neurons. J Neurosci 22:10277–10290.

Blair NT, Bean BP (2003) Role of tetrodotoxin-resistant Na+ current slow inactivation in adaptation of action potential firing in small-diameter dorsal root ganglion neurons. J Neurosci 23:10338–10350.

Bouheraoua N, Fouquet S, Marcos-Almaraz MT, Karagogeos D, Laroche L, Chédotal A (2019) Genetic Analysis of the Organization, Development, and Plasticity of Corneal Innervation in Mice. J Neurosci 39:1150–1168.

Browne LE, Latremoliere A, Lehnert BP, Grantham A, Ward C, Alexandre C, Costigan M, Michoud F, Roberson DP, Ginty DD, Woolf CJ (2017) Time-Resolved Fast Mammalian Behavior Reveals the Complexity of Protective Pain Responses. Cell Rep 20:89–98.

Cain DM, Wacnik PW, Turner M, Wendelschafer-Crabb G, Kennedy WR, Wilcox GL, Simone DA (2001) Functional interactions between tumor and peripheral nerve: changes in excitability and morphology of primary afferent fibers in a murine model of cancer pain. J Neurosci 21:9367–9376.

Chartier SR, Mitchell SAT, Majuta LA, Mantyh PW (2018) The Changing Sensory and Sympathetic Innervation of the Young, Adult and Aging Mouse Femur. Neuroscience 387:178–190.

Chiang H, Chang KC, Kan HW, Wu SW, Tseng MT, Hsueh HW, Lin YH, Chao CC, Hsieh ST (2018) Physiological and pathological characterization of capsaicin-induced reversible nerve degeneration and hyperalgesia. Eur J Pain 22:1043–1056.

Cho K, Jang JH, Kim SP, Lee SH, Chung SC, Kim IY, Jang DP, Jung SJ (2016) Analysis of Nociceptive Information Encoded in the Temporal Discharge Patterns of Cutaneous C-Fibers. Front Comput Neurosci 10.

Cummins TR, Dib-Hajj SD, Black JA, Akopian AN, Wood JN, Waxman SG (1999) A novel persistent tetrodotoxin-resistant sodium current in SNS-null and wild-type small primary sensory neurons. J Neurosci 19:RC43.

Dezhdar T, Moshourab RA, Fründ I, Lewin GR, Schmuker M (2015) A Probabilistic Model for Estimating the Depth and Threshold Temperature of C-fiber Nociceptors. In: Sci Rep.

Dib-Hajj SD, Cummins TR, Black JA, Waxman SG (2010) Sodium channels in normal and pathological pain. Annu Rev Neurosci 33:325–347.

Dubin AE, Patapoutian A (2010) Nociceptors: the sensors of the pain pathway. J Clin Invest 120:3760–3772.

Duraku LS, Hossaini M, Hoendervangers S, Falke LL, Kambiz S, Mudera VC, Holstege JC, Walbeehm ET, Ruigrok TJ (2012) Spatiotemporal dynamics of re-innervation and hyperinnervation patterns by uninjured CGRP fibers in the rat foot sole epidermis after nerve injury. Mol Pain 8:61.

Ferrante M, Migliore M, Ascoli GA (2013) Functional impact of dendritic branch-point morphology. J Neurosci 33:2156–2165.

Gemes G, Koopmeiners A, Rigaud M, Lirk P, Sapunar D, Bangaru ML, Vilceanu D, Garrison SR, Ljubkovic M, Mueller SJ, Stucky CL, Hogan QH (2013) Failure of action potential propagation in sensory neurons: mechanisms and loss of afferent filtering in C-type units after painful nerve injury. J Physiol 591:1111–1131.

Gidon A, Segev I (2012) Principles governing the operation of synaptic inhibition in dendrites. Neuron 75:330–341.

Gold MS, Gebhart GF (2010) Nociceptor sensitization in pain pathogenesis. Nat Med 16:1248–1257.

Goldstein RH, Katz B, Lev S, Binshtok AM (2017) Ultrafast optical recording reveals distinct capsaicin-induced ion dynamics along single nociceptive neurite terminals in vitro. J Biomed Opt 22:76010.

Goldstein RH, Barkai O, Íñigo-Portugués A, Katz B, Lev S, Binshtok AM (2019) Location and Plasticity of the Sodium Spike Initiation Zone in Nociceptive Terminals In Vivo. Neuron 102:801–812.

Grubb MS, Shu Y, Kuba H, Rasband MN, Wimmer VC, Bender KJ (2011) Short-and long-term plasticity at the axon initial segment. J Neurosci 31:16049–16055.

Grunditz A, Holbro N, Tian L, Zuo Y, Oertner TG (2008) Spine neck plasticity controls postsynaptic calcium signals through electrical compartmentalization. J Neurosci 28:13457–13466.

Gudes S, Barkai O, Caspi Y, Katz B, Lev S, Binshtok AM (2015) The role of slow and persistent TTX-resistant sodium currents in acute tumor necrosis factor-alpha-mediated increase in nociceptors excitability. J Neurophysiol 113:601–619.

Gulledge AT, Bravo JJ (2016) Neuron Morphology Influences Axon Initial Segment Plasticity. eNeuro 3:0085–0015.

Heppelmann B, Messlinger K, Neiss WF, Schmidt RF (1994) Mitochondria in fine afferent nerve fibres of the knee joint in the cat: a quantitative electron-microscopical examination. Cell Tissue Res 275:493–501.

Heppelmann B, Gallar J, Trost B, Schmidt RF, Belmonte C (2001) Three-dimensional reconstruction of scleral cold thermoreceptors of the cat eye. J Comp Neurol 441:148–154.

Herzog RI, Cummins TR, Waxman SG (2001) Persistent TTX-resistant Na+ current affects resting potential and response to depolarization in simulated spinal sensory neurons. J Neurophysiol 86:1351–1364.

Ivanusic JJ, Wood RJ, Brock JA (2013) Sensory and sympathetic innervation of the mouse and guinea pig corneal epithelium. J Comp Neurol 521:877–893.

Jadi M, Polsky A, Schiller J, Mel BW (2012) Location-dependent effects of inhibition on local spiking in pyramidal neuron dendrites. PLoS Comput Biol 8:14.

Jarvis S, Nikolic K, Schultz SR (2018) Neuronal gain modulability is determined by dendritic morphology: A computational optogenetic study. PLoS Comput Biol 14.

Johansson RS, Birznieks I (2004) First spikes in ensembles of human tactile afferents code complex spatial fingertip events. Nat Neurosci 7:170–177.

Kispersky TJ, Caplan JS, Marder E (2012) Increase in sodium conductance decreases firing rate and gain in model neurons. J Neurosci 32:10995–11004.

Kleggetveit IP, Namer B, Schmidt R, Helas T, Ruckel M, Orstavik K, Schmelz M, Jorum E (2012) High spontaneous activity of C-nociceptors in painful polyneuropathy. Pain 153:2040–2047.

Komagiri Y, Kitamura N (2003) Effect of intracellular dialysis of ATP on the hyperpolarization-activated cation current in rat dorsal root ganglion neurons. J Neurophysiol 90:2115–2122.

Kuba H, Ishii TM, Ohmori H (2006) Axonal site of spike initiation enhances auditory coincidence detection. Nature 444:1069–1072.

Kuba H, Oichi Y, Ohmori H (2010) Presynaptic activity regulates Na(+) channel distribution at the axon initial segment. Nature 465:1075–1078.

Lakatos S, Jancsó G, Horváth Á, Dobos I, Sántha P (2020) Longitudinal Study of Functional Reinnervation of the Denervated Skin by Collateral Sprouting of Peptidergic Nociceptive Nerves Utilizing Laser Doppler Imaging. Front Physiol 11.

Leibovich H, Buzaglo N, Tsuriel S, Peretz L, Caspi Y, Katz B, Lev S, Lichtstein D, Binshtok AM (2020) Abnormal Reinnervation of Denervated Areas Following Nerve Injury Facilitates Neuropathic Pain. Cells 9.

Lesniak DR, Marshall KL, Wellnitz SA, Jenkins BA, Baba Y, Rasband MN, Gerling GJ, Lumpkin EA (2014) Computation identifies structural features that govern neuronal firing properties in slowly adapting touch receptors. Elife 3:21.

Levy D, Tal M, Höke A, Zochodne DW (2000) Transient action of the endothelial constitutive nitric oxide synthase (ecNOS) mediates the development of thermal hypersensitivity following peripheral nerve injury. Eur J Neurosci 12:2323–2332.

Liu CN, Michaelis M, Amir R, Devor M (2000) Spinal nerve injury enhances subthreshold membrane potential oscillations in DRG neurons: relation to neuropathic pain. J Neurophysiol 84:205–215.

Magee JC (2000) Dendritic integration of excitatory synaptic input. Nat Rev Neurosci 1:181–190.

Magee JC, Johnston D (2005) Plasticity of dendritic function. Curr Opin Neurobiol 15:334–342.

Mainen ZF, Sejnowski TJ (1996) Influence of dendritic structure on firing pattern in model neocortical neurons. Nature 382:363–366.

Mancini F, Bauleo A, Cole J, Lui F, Porro CA, Haggard P, Iannetti GD (2014) Whole-body mapping of spatial acuity for pain and touch. Ann Neurol 75:917–924.

Manor Y, Koch C, Segev I (1991) Effect of geometrical irregularities on propagation delay in axonal trees. Biophys J 60:1424–1437.

Miyasho T, Takagi H, Suzuki H, Watanabe S, Inoue M, Kudo Y, Miyakawa H (2001) Low-threshold potassium channels and a low-threshold calcium channel regulate Ca2+ spike firing in the dendrites of cerebellar Purkinje neurons: a modeling study. Brain Res 891:106–115.

Murthy SE, Loud MC, Daou I, Marshall KL, Schwaller F, Kühnemund J, Francisco AG, Keenan WT, Dubin AE, Lewin GR, Patapoutian A (2018) The mechanosensitive ion channel Piezo2 mediates sensitivity to mechanical pain in mice. Sci Transl Med 10.

Müller LJ, Pels L, Vrensen GF (1996) Ultrastructural organization of human corneal nerves. Invest Ophthalmol Vis Sci 37:476–488.

Nimchinsky EA, Sabatini BL, Svoboda K (2002) Structure and function of dendritic spines. Annu Rev Physiol 64:313–353.

Nita, II, Caspi Y, Gudes S, Fishman D, Lev S, Hersfinkel M, Sekler I, Binshtok AM (2016) Privileged crosstalk between TRPV1 channels and mitochondrial calcium shuttling machinery controls nociception. Biochim Biophys Acta 1863:2868–2880.

Ofer N, Shefi O, Yaari G (2017) Branching morphology determines signal propagation dynamics in neurons. Sci Rep 7:8877.

Olivares E, Salgado S, Maidana JP, Herrera G, Campos M, Madrid R, Orio P (2015) TRPM8-Dependent Dynamic Response in a Mathematical Model of Cold Thermoreceptor. PLoS One 10:e0139314.

Olson W, Abdus-Saboor I, Cui L, Burdge J, Raabe T, Ma M, Luo W (2017) Sparse genetic tracing reveals regionally specific functional organization of mammalian nociceptors. Elife 12:29507.

Panzeri S, Petersen RS, Schultz SR, Lebedev M, Diamond ME (2001) The role of spike timing in the coding of stimulus location in rat somatosensory cortex. Neuron 29:769–777.

Persson AK, Black JA, Gasser A, Cheng X, Fischer TZ, Waxman SG (2010) Sodium-calcium exchanger and multiple sodium channel isoforms in intra-epidermal nerve terminals. Mol Pain 6:84.

Qu L, Caterina MJ (2016) Enhanced excitability and suppression of A-type K+ currents in joint sensory neurons in a murine model of antigen-induced arthritis. In: Sci Rep.

Reeh PW (1986) Sensory receptors in mammalian skin in an in vitro preparation. Neurosci Lett 66:141–146.

Rho YA, Prescott SA (2012) Identification of molecular pathologies sufficient to cause neuropathic excitability in primary somatosensory afferents using dynamical systems theory. PLoS Comput Biol 8:24.

Sandkuhler J (2009) Models and mechanisms of hyperalgesia and allodynia. Physiological reviews 89:707–758.

Segev I, Rall W (1988) Computational study of an excitable dendritic spine. J Neurophysiol 60:499–523.

Shah MM, Migliore M, Valencia I, Cooper EC, Brown DA (2008) Functional significance of axonal Kv7 channels in hippocampal pyramidal neurons. Proc Natl Acad Sci U S A 105:7869–7874.

Sjöström PJ, Rancz EA, Roth A, Häusser M (2008) Dendritic excitability and synaptic plasticity. Physiol Rev 88:769–840.

St Pierre M, Reeh PW, Zimmermann K (2009) Differential effects of TRPV channel block on polymodal activation of rat cutaneous nociceptors in vitro. Experimental brain research Experimentelle Hirnforschung Expérimentation cérébrale 196:31–44.

Sundt D, Gamper N, Jaffe DB (2015) Spike propagation through the dorsal root ganglia in an unmyelinated sensory neuron: a modeling study. J Neurophysiol 114:3140–3153.

Tal M, Devor M (1992) Ectopic discharge in injured nerves: comparison of trigeminal and somatic afferents. Brain Res 579:148–151.

Treede RD, Meyer RA, Campbell JN (1990) Comparison of heat and mechanical receptive fields of cutaneous C-fiber nociceptors in monkey. J Neurophysiol 64:1502–1513.

Vandewauw I, De Clercq K, Mulier M, Held K, Pinto S, Van Ranst N, Segal A, Voet T, Vennekens R, Zimmermann K, Vriens J, Voets T (2018) A TRP channel trio mediates acute noxious heat sensing. Nature 555:662–666.

Vasylyev DV, Waxman SG (2012) Membrane properties and electrogenesis in the distal axons of small dorsal root ganglion neurons in vitro. J Neurophysiol 108:729–740.

Waxman SG, Zamponi GW (2014) Regulating excitability of peripheral afferents: emerging ion channel targets. Nat Neurosci 17:153–163.

Weidner C, Schmidt R, Schmelz M, Torebjork HE, Handwerker HO (2003) Action potential conduction in the terminal arborisation of nociceptive C-fibre afferents. J Physiol 547:931–940.

Williams SR, Stuart GJ (2003) Role of dendritic synapse location in the control of action potential output. Trends Neurosci 26:147–154.

Woolf CJ, Ma Q (2007) Nociceptors--noxious stimulus detectors. Neuron 55:353–364.

Yamada R, Kuba H (2016) Structural and Functional Plasticity at the Axon Initial Segment. Front Cell Neurosci 10.

Zimmermann K, Hein A, Hager U, Kaczmarek JS, Turnquist BP, Clapham DE, Reeh PW (2009) Phenotyping sensory nerve endings in vitro in the mouse. Nature protocols 4:174–196.

Zylka MJ, Rice FL, Anderson DJ (2005) Topographically distinct epidermal nociceptive circuits revealed by axonal tracers targeted to Mrgprd. Neuron 45:17–25.

